# Exaggerated male hindlegs function as pure weapons of male–male combat in thorny devil stick insects

**DOI:** 10.1101/2025.11.21.689692

**Authors:** Romain P. Boisseau, Douglas J. Emlen

**Author notes:** Corresponding author: R.P.B.

## Abstract

Sexually selected weapons can function as both combat tools and agonistic signals of fighting ability, depending on whether and how males assess rivals. We investigated the function of the enlarged male hindlegs in the New Guinean thorny devil stick insects, *Eurycantha calcarata* in male-male and male-female interactions. Field and lab experiments showed that larger males with proportionally larger hindlegs were more likely to win fights over access to females and subsequently mate. Behavioral sequence analyses and contest cost predictors indicated that males likely use a mutual assessment strategy. Surprisingly, males did not appear to use their hindlegs as signals of fighting ability, relying instead on tactile and chemical cues to assess opponents. Hindlegs were employed only to deliver powerful squeezes in rare, escalated fights. During copulation, males also used hindlegs to stabilize their position, but as females did not appear to resist, we found no evidence for a coercive function. These findings suggest that enlarged male hindlegs in *E. calcarata* serve purely as force-delivering combat tools rather than signaling structures, even though males assess rivals during contests. These result highlight how understanding the specific functions and contexts of weapon use provides critical insight into the diversification of sexually selected traits.

**Teaser text:** Sexually selected weapons vary widely in function, serving as signals, combat tools, or both—which shapes their evolution. In the thorny devil stick insect (*Eurycantha calcarata*), males use enlarged, spined hindlegs solely as force-delivering weapons in direct combat with rivals, not as displays or coercive tools. Males assess opponents using tactile or chemical cues, but their hindlegs scale proportionately with body size, reflecting selection for mechanical performance rather than signaling. These findings highlight that understanding the behavioral context and function of weapon use is essential to explaining their evolutionary diversification.

## 1. Introduction

Sexual selection often drives the evolution of exaggerated, sexually dimorphic traits, including weapons used in male–male competition for mates or resources (Andersson 1994; Emlen 2008; Shuker and Simmons 2014). Intra-sexually selected weapons are first and foremost tools of battle between same-sex rivals (Rico-Guevara and Hurme 2019), but they often serve additional roles. For instance, enlarged claws of male fiddler crabs deter predators (Bildstein et al. 1989; McLain et al. 2003), while antlers in male deer are also used to harass females (Clutton-Brock and Parker 1995; Pradhan and Van Schaik 2009). More commonly, weapons double as visual or tactile signals of quality that threaten rivals or attract females (Berglund et al. 1996; Pratt et al. 2003; Muramatsu 2011; Rometsch et al. 2021). Such traits can be thought of as falling along a continuum between weapon and signal (McCullough et al. 2016), depending on the relative importance of biomechanical performance and conspicuousness (McCullough and O’Brien 2022). Predominantly signaling weapons tend to exhibit exaggerated elaboration and positive static allometry, amplifying size differences and aiding opponent assessment (Eberhard et al. 2018; O’Brien et al. 2018; Rodríguez and Eberhard 2019). In contrast, purely force-delivering weapons are constrained to maintain lever proportions and avoid the “paradox of the weakening combatant,” where larger weapons lose efficiency (Levinton and Allen 2005; O’Brien and Boisseau 2018). Therefore, the functional details of weapons and the different contexts in which they are used critically influence their evolution.

Understanding how weapons are used in combat requires understanding how animals decide when to initiate, escalate, or retreat from fights, how long to persist, and how much cost to endure (Hardy and Briffa 2013). Contest theory, grounded in game theory, investigates factors affecting optimal choices made in the presence of other decision-makers (Maynard Smith and Price 1973; Maynard Smith 1974, 1982). The loser ultimately decides when to withdraw, based on factors such as resource value (Maynard Smith and Parker 1976; Arnott and Elwood 2008), prior fighting experience (Hsu et al. 2006; Rutte et al. 2006; Goubault and Decuignière 2012), and relative fighting ability or resource-holding potential (RHP) (Parker 1974; Maynard Smith and Parker 1976; Arnott and Elwood 2009). Theoretical studies describe two main assessment strategies of RHP: self-assessment and mutual assessment (Taylor and Elwood 2003; Arnott and Elwood 2009; Chapin et al. 2019). Self-assessment models encompass both pure self-assessment and cumulative assessment models (CAM). In pure self-assessment models, contestants monitor only their own energetic state and cease fighting when internal cost thresholds are reached, regardless of their opponent’s actions (Mesterton-Gibbons et al. 1996; Payne and Pagel 1996; Arnott and Elwood 2009). In contrast, CAM allows costs to be imposed by the opponent (e.g., through injury or energy loss), so the weaker individual gives up first because it accumulates costs faster (Payne 1998; Arnott and Elwood 2009). Under mutual assessment, contestants estimate both their own and their rival’s RHP, and fights end when one recognizes its inferiority (Enquist and Leimar 1983). These models yield distinct predictions that have been widely tested (Arnott and Elwood 2009; Elwood and Arnott 2012; Green and Patek 2018; Chapin et al. 2019; Pinto et al. 2019) (Fig. 1). Mutual assessment offers the advantage of early contest resolution and reduced costs, yet empirical studies more often support self-assessment strategies (Elwood and Arnott 2013; Pinto et al. 2019).

**Figure 1:**
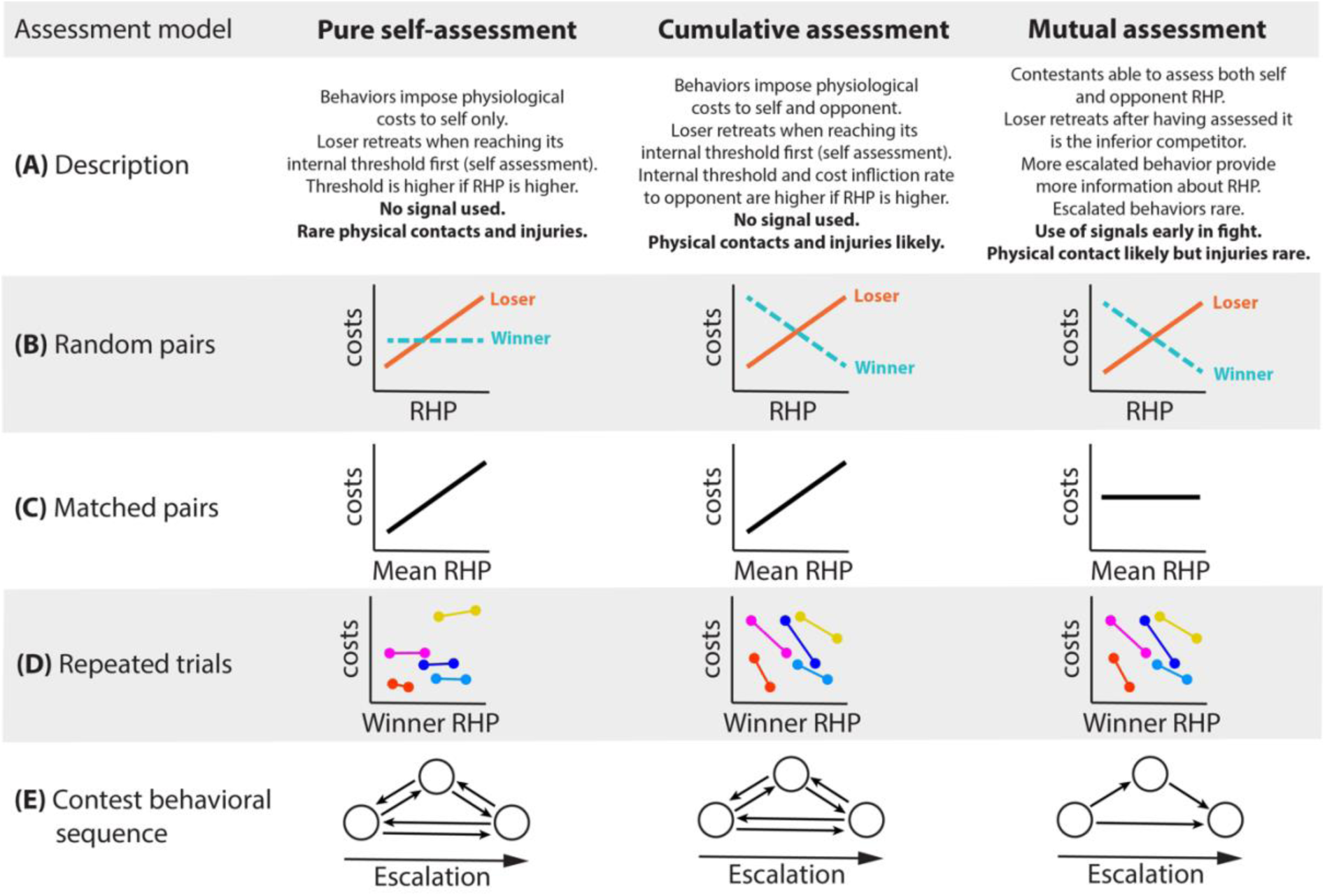
Main assessment models and associated predictions. **(A)** The rationale of each model is described; predictions related to the likelihood of signaling, physical contact, and injuries are indicated in bold. (**B**) Models predict different relationships between contest costs and RHP for losers (orange line) and winners (turquoise line) of randomly matched contests, and (**C**) for the averaged RHP of contestants in matched contests (black line) (Arnott and Elwood 2009; Green and Patek 2018). (**D**) For trials where focal animals with a low RHP (i.e., losers) are assigned to multiple opponents with higher RHP (i.e., winners), models also predict different relationships between winner RHP and contest cost for each focal animal (Chapin et al. 2019). For each focal animal (represented by a different color), a negative slope indicates that winner RHP affects contest cost, which is only expected under the cumulative or mutual assessment models. No relationship is expected under a pure self-assessment strategy. This approach is used to discern heterogeneity in the strategy used by individuals within a population. Finally, (**E**) models predict different trends in the directionality with which contest behaviors unfold (circles correspond to behaviors, arrows represent most likely transitions between them) (Green and Patek 2018).

Integrating studies of function, allometry, and contest assessment strategies is essential for understanding the evolution of sexually selected weapons. For instance, positively allometric weapons that serve as threat signals are expected only in systems where males assess their rivals’ RHP during combat – that is, in species with mutual assessment. If contest escalation decisions are made based on internal assessments of energetic state or damage alone, there would be no selection for hypervariability in weapon expression since there would be no value to receivers in signal traits that amplify differences in RHP. Instead, selection should favor weapon allometries that preserve biomechanical performance across a range of body sizes. Here, we examine the fighting behavior and weapon function of male New Guinean thorny devil stick insects (*Eurycantha calcarata*, Lucas, 1869; Lonchodinae, Phasmatodea) using laboratory and field data. In this nocturnal species, males are relatively large compared to other phasmid species, and possess greatly enlarged hind femora armed with a sharp spine (Buckley et al. 2009). Field observations revealed that this species displays a territory- and female-defense mating system where males fight with rivals, notably using their hindlegs, for strategic positions close to tree cavities where adults communally roost during the day. Territory holders are then able to intercept females as they leave their shelter at dusk, and copulate (Boisseau et al. 2020). Puncture wounds on male hind femora attest to the dangerous nature of these weapons (Boisseau et al. 2020; Lane and McCullough 2025). Although field data clearly indicate a combat role for the hindlegs, quantitative links between body or weapon size and fighting success remain untested. Males also use their hindlegs defensively against predators, raising them in a threat posture and striking when approached (Bedford 1976; Carlberg 1989; Boisseau et al. 2020; Koczur et al. 2024). Finally, males wrap one hindleg around the female’s abdomen during copulation (Hsiung 1987), suggesting an additional role in mating coercion, as seen in other insects with sexually dimorphic hindlegs (Rowe et al. 2006; Haley and Gray 2012; Burrows 2020).

In this study, we tested whether male *E. calcarata* hindlegs function as pure weapons, threat signals, or coercive tools. We examined how body and weapon size affect fight outcomes and the relationship between fighting and mating success. We also evaluated different contest assessment models to determine how males gauge rivals’ RHP and whether hindlegs serve as signals. Finally, we investigated whether larger hindlegs aid males in coercing females and whether females benefit from resisting copulation to reproduce via parthenogenesis, as suggested in other phasmids (Burke et al. 2015; Burke and Bonduriansky 2022).

## 2. Materials and methods

Statistical analyses are detailed in each relevant section of the Materials and Methods. All analyses in this study were performed using R v4.1.1 (R Core Team 2023). Linear models were checked for normality of residuals and absence of patterning. R functions and packages are indicated throughout the materials and methods as *function*: ‘package’.

### Study animals and measurements

We used a culture population of *E. calcarata* originally collected around Kimbe (West New Britain, PNG) in the late 1970s. The insects were housed in transparent plastic containers (65 × 45 × 50cm) at 22°C, 12h:12h light:dark, 50–80% relative humidity and fed *ad libitum* on maple leaves (*Acer platanoides*). Males and females were reared together (∼50 per container) until adulthood, then separated.

Hindleg ontogeny was measured on 47 males and 66 females at four developmental stages—fourth, fifth, sixth instars, and adult—using photographs (Canon EOS 600D) and ImageJ (Schneider et al. 2012). Traits included mesothorax length —used as a proxy for body size (Boisseau et al. 2020)—right front femur length, right hind femur length, and width. Hind femur area was calculated as an ellipse 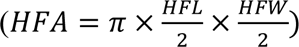 and validated against manually outlined adult femurs (Fig. S1). To investigate changes in scaling relationships between body size and front or hind leg size during postembryonic development, we ran linear mixed models (LMM, *lme*: ‘nlme’, Pinheiro et al. 2021) using either log_10_-transformed leg measurements as response variable and log_10_ mesothorax length, sex and instar as predictor variables as well as all two-way and three-way interactions. Individual ID was added as a random factor. A sequential ANOVA (*anova.lme*: ‘nlme’) was used to assess the significance of the fixed effects. Non-significant interaction terms (p>0.05) were deleted from the model to provide the most accurate parameter estimates. Adult scaling relationships were assessed for deviations from isometry (slope = 1 for linear traits, slope = 2 for surface traits) using 95% confidence intervals.

### Field observations

Field observations were conducted near Kimbe, Papua New Guinea (Dami palm plantations, S5° 31.846′ E150° 20.221′) and are described more fully in Boisseau et al. (2020). Adult male behaviors around tree cavities were recorded during four full nights on a *Kleinhovia hospita* trunk with 12 cavities. Time-lapse video cameras (HERO4, GoPro; 0.5 s interval) recorded interactions under red light from 4 PM to 8 AM.

Fights were defined as contact between two males resulting in one male retreating from the fighting area (loser) and one maintaining position (winner). 33 fights among 31 male pairs were observed over females or territories near cavity entrances. One male was randomly assigned as focal and the other as opponent. Fight outcome was recorded as a binary variable: 1 if the focal male won by retaining control of the resource or forcing the opponent to retreat, and 0 if he lost by retreating. Ownership was assigned to the male that either already occupied the territory or was already with the female. BORIS v7.5.3 (Friard and Gamba, 2016) was used to code behaviors (Table 1). Although some males had been measured and marked prior to observations, many individuals involved in agonistic interactions were unmarked and could not be physically measured. Consequently, absolute body size was often unavailable, and we estimated relative size differences between contestants by measuring their body lengths directly from video recordings.

**Table 1:** Ethogram and description of the recorded fighting behaviors.

| Behavior | Description |
| --- | --- |
| Start | Beginning of the fight: first interaction between the two contestants. |
| Antennate | The contestant touches its opponent (often repeatedly) with its antennae. |
| Kick | The contestant briefly kicks its opponent with either its front, middle or hind legs. |
| Push | The contestant lunges towards its opponent |
| Chase | The contestant chases its opponent who is running away |
| Mount | The contestant climbs on top of its opponent |
| Descend | The contestant dismounts its opponent |
| Turn | The contestant does a 180° rotation (often while on top of its opponent) |
| Back | The contestant moves backward towards its opponent |
| Grasp | The contestant grabs and squeezes its opponent with its hindlegs (often while on top of it) |
| Knock | The contestant repeatedly hammers the substrate with the end of its abdomen |
| End | End of the fight: the loser retreats away from the winner. |

### Laboratory fighting trials

#### Arena setup

Trials took place in glass arenas (90 × 45 × 45 cm) whose bottom was covered with moist paper, dried maple leaves, two water dishes, and an artificial roosting cavity (35 × 16 cm) consisting of maple bark with a 4 cm hole and cardboard rim to restrict side access. White fluorescent lights provided daytime illumination (12h), red lights illuminated nighttime periods. The inside of the cavity was constantly lit in red light. Two cameras (HERO4, GoPro, San Mateo, CA, USA; interval: 0.5s) recorded inside and outside the cavity (Fig. S2).

#### Randomly-matched trials

To investigate how size affected fighting and mating success, we staged trials involving three randomly picked adult males and three adult females (lab-bred). Individuals were weighed before and after trials (ME104T/00, Mettler Toledo, Columbus, OH, USA) and observed over four consecutive days. 17 such trials could be run involving 22 different males and 17 females. Thus, most individuals were used in several trials. Individuals (males and females) were involved in a trial approximately every month (±1week), starting two weeks after their final molt and until they died. Trials always involved different combinations of individuals. Some females involved in these trials, all of whom mated during the trials, were individually followed to record their fecundity (n=12, see section *Effect of mating on female fitness*). For every male-male fight outside the cavity, we recorded the outcome, ownership and the detailed sequence of agonistic behaviors (Video S1).

We also recorded male interactions inside the cavity. During the daytime, one male would typically sit on or close to the females and exclude the other males to the opposite corner away from the females (Video S2). Individual contests were difficult to delineate as losers could not clearly retreat away from the winner. Thus, we considered fights to last the entire time two males were found together in the cavity with at least one female. The male sitting the closest to the females and clearly successfully guarding them was considered the winner. Day periods for which only one male was present in the cavity or for which the winner was unclear were not considered.

#### Body size-matched trials

To be able to test the effect of relative weapon size on fight outcome and distinguish between the cumulative and mutual assessment models (Fig. 1D), we used another set of lab-bred individuals to perform body size-matched contests. During these trials (n=21), only two males and one female were simultaneously introduced and filmed in the arena for three days. Pairs of males were matched by mesothorax length (i.e., body size) and did not differ by more than 5%. We used 40 different males, among which two were involved in two trials, and 21 different 2-weeks old virgin females. In other respects, we followed the same procedures detailed above.

### Behavioral scoring and fight outcome analyses

For all field, randomly matched, and size-matched lab fights, we calculated contestant size asymmetry using the following index:

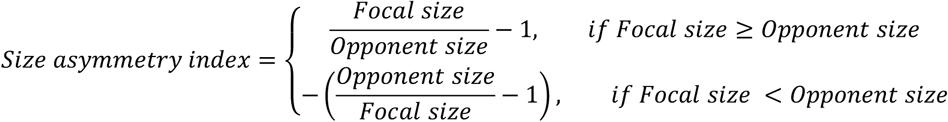

Size corresponded to body length for field contests, and to mesothorax length, body mass, and hind femur area for lab contests. This index, analogous to the sexual dimorphism index of Lovich and Gibbons (1992), is symmetrical and centered on zero regardless of which contestant is larger.

For field contests, we tested the effect of body length asymmetry on contest outcome using a generalized linear mixed-effects model (GLMM; binomial family, logit link; *glmer*, “lme4”). The asymmetry index was scaled to unit variance and included as a fixed effect with ownership. Male pair ID was included as a random intercept to account for repeated fights between the same pairs, though pseudoreplication due to individuals appearing in multiple fights could not be controlled. Significance of fixed effects was assessed using chi-square tests (*anova*, “stats”).

For lab contests, asymmetry indexes were calculated using mesothorax length, mean body mass (before and after trials), and hind femur area. We observed 140 randomly matched and 188 size-matched contests outside the roosting cavity (51 and 21 male pairs, respectively) and 73 randomly-matched contests inside (33 male pairs). Contest data were compiled in long format (sensu Briffa et al. 2013) with two rows per fight (one per contestant) to allow GLMMs accounting for contestant replication. We then tested the effects of body size, weapon size, body mass, and ownership on win probability using GLMMs (*glmer*, “lme4”), with contest outcome as the response and mesothorax length, hind femur area, body mass, and ownership as fixed effects; fight ID, contestant ID, and trial ID as random effects. All size variables were log10-transformed, scaled to unit variance, and centered prior to analysis. Fixed-effect significance was evaluated sequentially using chi-square tests (anova, “stats”).

### Sequential behavioral analyses

Assessment models predict that contest behaviors either escalate in distinct phases or occur randomly (Fig. 1E). To test how male fighting behaviors fit these predictions, we conducted a sequential analysis following Green and Patek (2018) using the R package *igraph* (Csardi and Nepusz 2006). Behavioral sequences from all field, randomly matched, and size-matched lab contests were combined separately, each summarized into an adjacency matrix where rows and columns represented the 12 contest behaviors (Table 1). Each matrix cell contained the frequency of transitions between behaviors, calculated by dividing transition counts by the total number of transitions.

To identify transitions occurring more often than expected by chance, we generated 10,000 randomized matrices preserving behavior frequencies but randomizing transitions. Transitions between “start” and fighting behaviors, “end” and fighting behaviors, or repeated “mount”/“descend” actions were disallowed. For each transition, we extracted the 95^th^ percentile of the null distribution and retained only observed transitions exceeding this threshold as significant.

Significant transitions were visualized as network graphs, where vertices represented behaviors (size proportional to behavior frequency) and edges represented significant transitions (width proportional to transition probability). These networks allowed us to detect contest phases—clusters of behaviors that frequently transition among themselves and rarely recur once a new phase begins (Enquist et al. 1990; Green and Patek 2018).

### Correlational tests of assessment models

Assessment models also differ in their predicted relationships between contestants’ RHP and contest cost (Fig. 1B–D), where cost is often indexed by contest duration and maximum escalation level (e.g., Fea and Holwell 2018; Green and Patek 2018). We used a composite contest cost metric integrating both duration and intensity. Each behavior was assigned an escalation score (1–5) based on its phase in the behavioral sequence: early and late low-intensity phases (antennal contacts, descending, substrate knocking) scored 1, and later phases with greater physical contact (kicking, pushing, chasing, mounting, hind-leg grasping) scored 2–5. Contest cost was the sum of all behavior scores, increasing with fight length and intensity.

We first examined the relationship between contest cost and RHP asymmetry, using mesothorax length as a proxy for RHP. Mutual assessment predicts a negative correlation, whereas self-assessment may produce similar patterns incidentally (Taylor and Elwood 2003). We combined data from randomly- and size-matched lab trials and fitted a linear mixed-effects model (LMM; lmer, “lme4”) with log_10_-transformed contest cost as the response and absolute mesothorax length asymmetry as a fixed effect; pair ID, trial ID, and year as random effects. Significance was tested via chi-square tests (*Anova*, “car”).

We next tested correlations between contest cost and loser or winner RHP in randomly matched contests (Fig. 1B), using separate LMMs with log₁₀ mesothorax length of either contestant as the fixed effect and loser (or winner) ID, pair ID, trial ID, and day (i.e., if the fight occurred on the first, second, third or fourth day of the trial) as random effects. We also modeled contest cost as a function of average contestant RHP for size-matched trials (Fig. 1C), with pair ID, trial ID, and day as random effects.

Finally, using our randomly-matched trials which involved three males and the repeated-trial approach of Chapin et al. (2019) (Fig. 1D), we assessed individual variation in assessment strategies. For each trial involving three males, the smallest male (focal loser) was used to test the effect of opponent RHP (winner mesothorax length) on contest cost. LMMs included log₁₀ contest cost as the response and log₁₀ opponent RHP, focal loser ID, and their interaction as fixed effects, with opponent ID and trial ID as random effects. Fixed-effect significance was tested sequentially with chi-square tests (*anova*, “stats”).

### Correlation between fighting and mating success

Using our first generation of lab insects (randomly matched trials with 3 males and 3 females), we tested whether males that won more fights also mated more often. For each male, we recorded absolute mating success (number of copulations) and calculated relative mating success by dividing it by the trial’s mean male mating success.

Fighting success was quantified as dominance rank within each male triad, based on overall contest outcomes across four days. A male dominating both rivals was ranked 1, an intermediate male 2, and the least dominant 3. In cases of ties (equal wins or no fights), both males received rank 1 if dominant over the third, or rank 2 if subordinate to it. Separate dominance ranks were first assigned for contests inside and outside the cavity, averaged, and rounded down to yield an overall rank used as a proxy for fighting success.

We then tested the relationship between fighting and mating success using a cumulative link mixed model (*clmm*, “ordinal”), with dominance rank (ordered factor) as the response, relative mating success as the fixed effect, and trial ID and male ID as random effects.

### Sexual conflict and male-female interactions

Using our first generation of lab insects (randomly matched trials with 3 males and 3 females), we examined male–female interactions to test whether males use their enlarged hindlegs coercively. For each mating, we recorded male latency to mate (time from first contact to copulation onset) and copulation duration. If hindlegs aid in coercion, larger males with larger hindlegs should initiate copulation faster and maintain longer copulations. We also considered female reproductive status (virgin vs. mated), as virgins may either accept mating more readily to fertilize eggs or resist to favor parthenogenesis. We analyzed latency to mate and copulation duration separately using linear mixed-effects models (LMMs), with male mesothorax length, male hind femur area, female mesothorax length, and female mating status as fixed effects, and male ID, female ID, and trial ID as random effects. Continuous variables were log₁₀-transformed, centered, and scaled to unit variance prior to analysis.

### Effect of mating on female fitness

To compare the fitness consequences of sexual versus parthenogenetic reproduction, we used females from our first generation. A subset of females (n = 16) was kept in female-only trials (“parthenogenetic” treatment), while others from the mixed trials served as “mated” females (n = 12). Female-only trials involved six females housed together for four days under conditions identical to the mixed trials. Each female participated roughly once a month, starting two weeks after the final molt and continuing until death.

Between trials, females were housed individually in small plastic containers (35 × 20 × 15cm) with a dirt box for oviposition, a water dish, and fed dried maple leaves *ad libitum*. Containers were cleaned weekly, and eggs were collected, counted, and weighed. Female survival was monitored daily. Eggs were kept on moist compost at 22 °C, with hatching rate and development time (from first egg laid to first hatching) recorded. Offspring were reared in similar containers in groups of 50 nymphs by treatment, maintained under identical conditions, and monitored for survival to second instar.

We compared adult female survival between treatments using survival analyses (*survdiff*, “survival”). Effects of treatment on fecundity (lifetime egg number, average egg mass, egg development time, egg-laying rate, hatching rate) were tested with linear models (*lm*, “stats”) including female mesothorax length and treatment as predictors. Continuous variables were log₁₀-transformed, and significance was assessed via Type I ANCOVA. Offspring survival to second instar was compared between treatments using a Fisher’s exact test (*fisher.test*, “stats”).

## 3. Results

### Ontogenetic scaling relationships

Males had relatively longer front femurs and longer, wider hind femurs than females, with these differences becoming more pronounced in the final two instars (Fig. 2, Table S1). In both sexes, relative front and hind femur dimensions increased through development. Allometric slopes did not differ between sexes, and adults showed slopes consistent with isometry for all measured traits (Fig. 2, Table S1). Hind femur length and width scaled isometrically, similar to front femur length, which served as an unspecialized reference trait (O’Brien et al. 2018). Thus, male hindleg exaggeration in *E. calcarata* results from an increased intercept between body and hindleg size late in development rather than from steeper allometric scaling.

**Figure 2:**
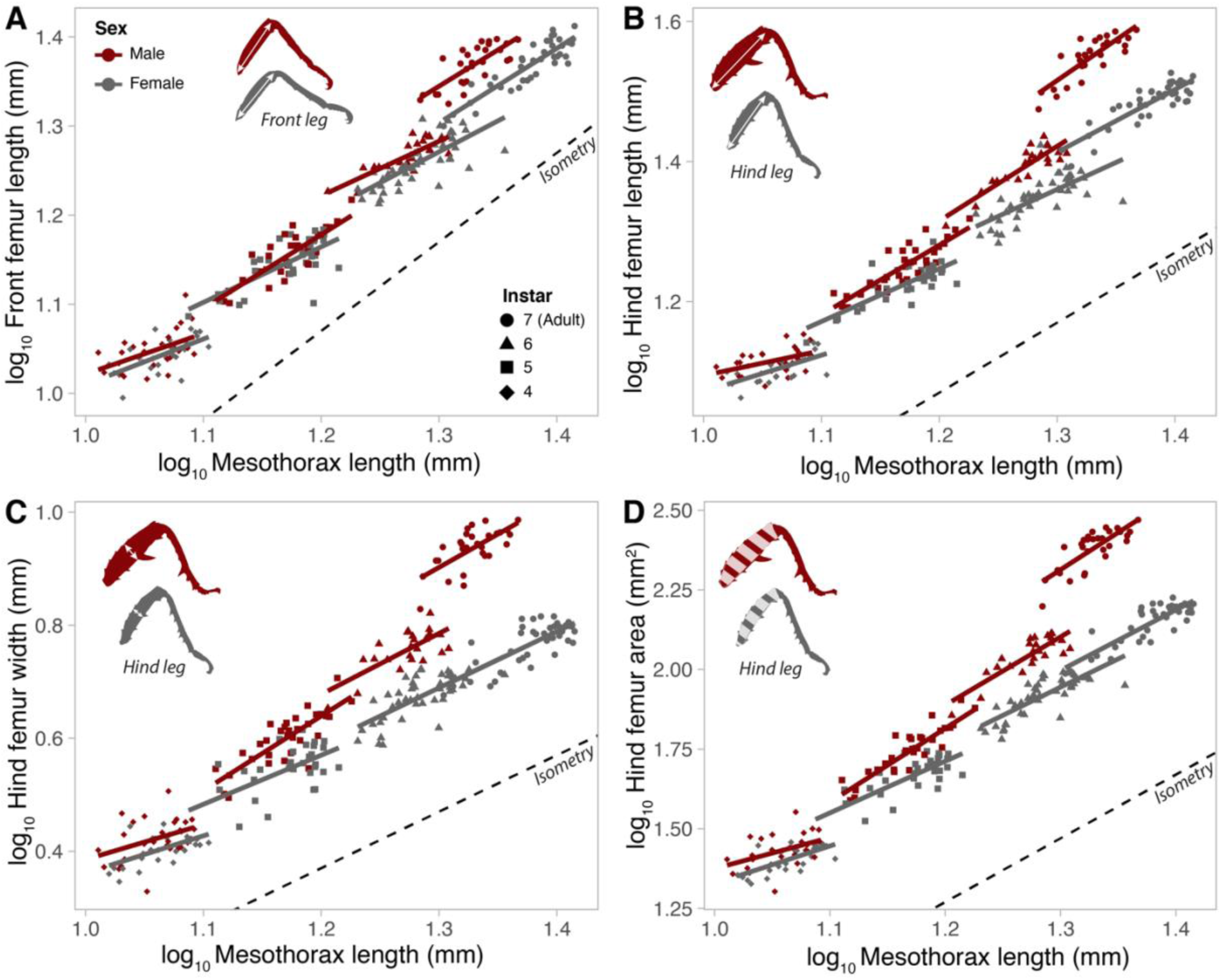
Sexual dimorphism and scaling relationships between body size and front or hind leg size across the second half of postembryonic development in lab-reared *E. calcarata*. Scaling relationships between front femur length (**A**), hind femur length (**B**), hind femur width (**C**) and hind femur area (**D**) and mesothorax length (∼body size) for males and females across the last three nymphal instars and adults. The dashed line represents an isometric slope (arbitrary intercept). Leg drawings in the top left corner illustrate the trait measured. Corresponding statistical analyses are reported in Table S1.

### Fight outcomes

In the field, both body length difference (χ^2^ = 17.3, df = 1, p < 0.0001) and ownership (χ^2^ = 7.01, df = 1, p = 0.008) significantly affected contest outcomes (Fig. 3A), while their interaction was non-significant (χ^2^ = 0.47, df = 1, p = 0.49). Larger individuals and residents were more likely to win.

**Figure 3:**
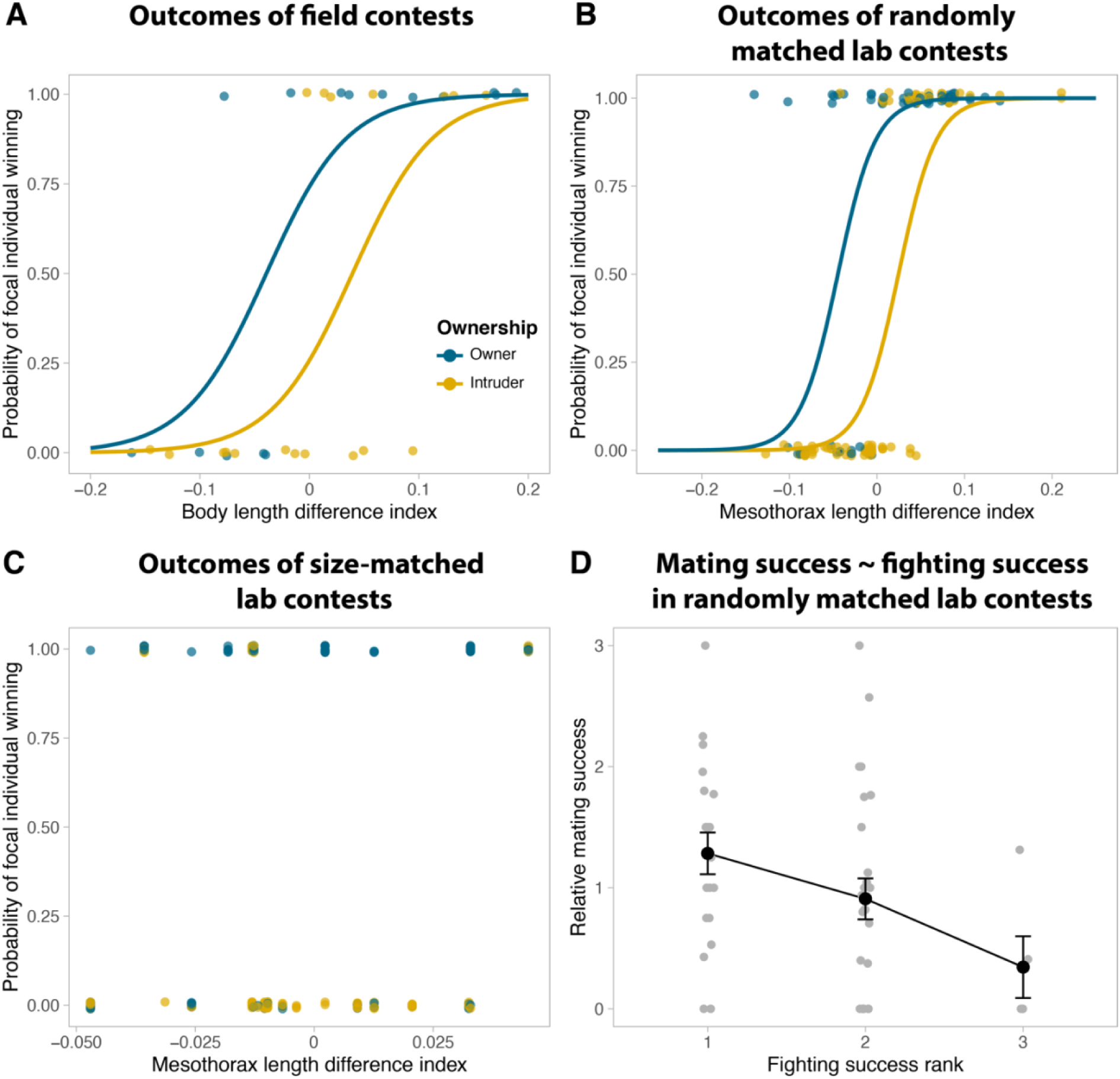
Male fight outcomes and correlation between fighting and mating success. Binary GLMM of (**A**) body length difference index in the field, mesothorax length difference index in randomly matched lab trials **(B)** and mesothorax length difference index in size-matched lab trials **(C)** against contest outcome. Colors represent the ownership status of the focal individual (resident/owner in blue, intruder in yellow). Correlation between relative mating success and male fighting rank in randomly-matched trials involving three males and three females. Black points correspond to means per fighting rank and error bars indicate standard errors to the mean.

In randomly matched lab contests, mesothorax length (χ^2^ = 18.24, df = 1, p < 0.0001) and ownership (χ^2^ = 40.03, df = 1, p < 0.0001) both predicted victory outside the roosting cavity, whereas hind femur area (χ^2^ = 0.42, p = 0.52) and body mass (χ^2^ = 0.12, p = 0.73) had no effect after accounting for body size (Fig. 3B). Similarly, mesothorax length positively affected the probability of winning contests inside a cavity (χ^2^= 5.93, df=1, p=0.015), but the effect of hind femur area (χ^2^= 2.33, df=1, p=0.13) and body mass (χ^2^= 1.06, df=1, p=0.30) were not significant after accounting for body size.

In size-matched lab contests, contest outcome depended only on ownership (χ² = 29.7, df = 1, p < 0.0001) (Fig. 3C). Mesothorax length (χ^2^= 0.004, df=1, p=0.95), hind femur area (χ^2^= 0.14, df=1, p=0.71) and body mass (χ^2^= 1.63, df=1, p=0.20) did not significantly affect contest outcome. Thus, residents consistently outperformed intruders, and the effects of body condition or weapon size differences were undetectable.

Finally, fighting rank correlated with relative mating success (z = −2.10, p = 0.035; Fig. 3D): dominant males that won more fights were more likely to mate during the four-day trials.

### Sequential behavioral analyses

Behavioral sequence analyses revealed that fights progressed through distinct escalation phases consistent with mutual assessment predictions (Fig. 1E, 4). In both field and lab contests, fights typically began with (1) antennal contact, followed by (2) brief kicks or pushes, leading either to (3) a chase or (4) mounting, and sometimes (5) grasping with the weaponized hindlegs (Fig. 4A). Fights then deescalated to (6) termination, often marked by the winner thumping its abdomen on the substrate while the loser retreated (Video S3). Within phases, behaviors occurred at similar frequencies, and no transitions returned to earlier phases (Fig. 4B–D). Lower-intensity behaviors were far more frequent than highly escalated ones.

**Figure 4:**
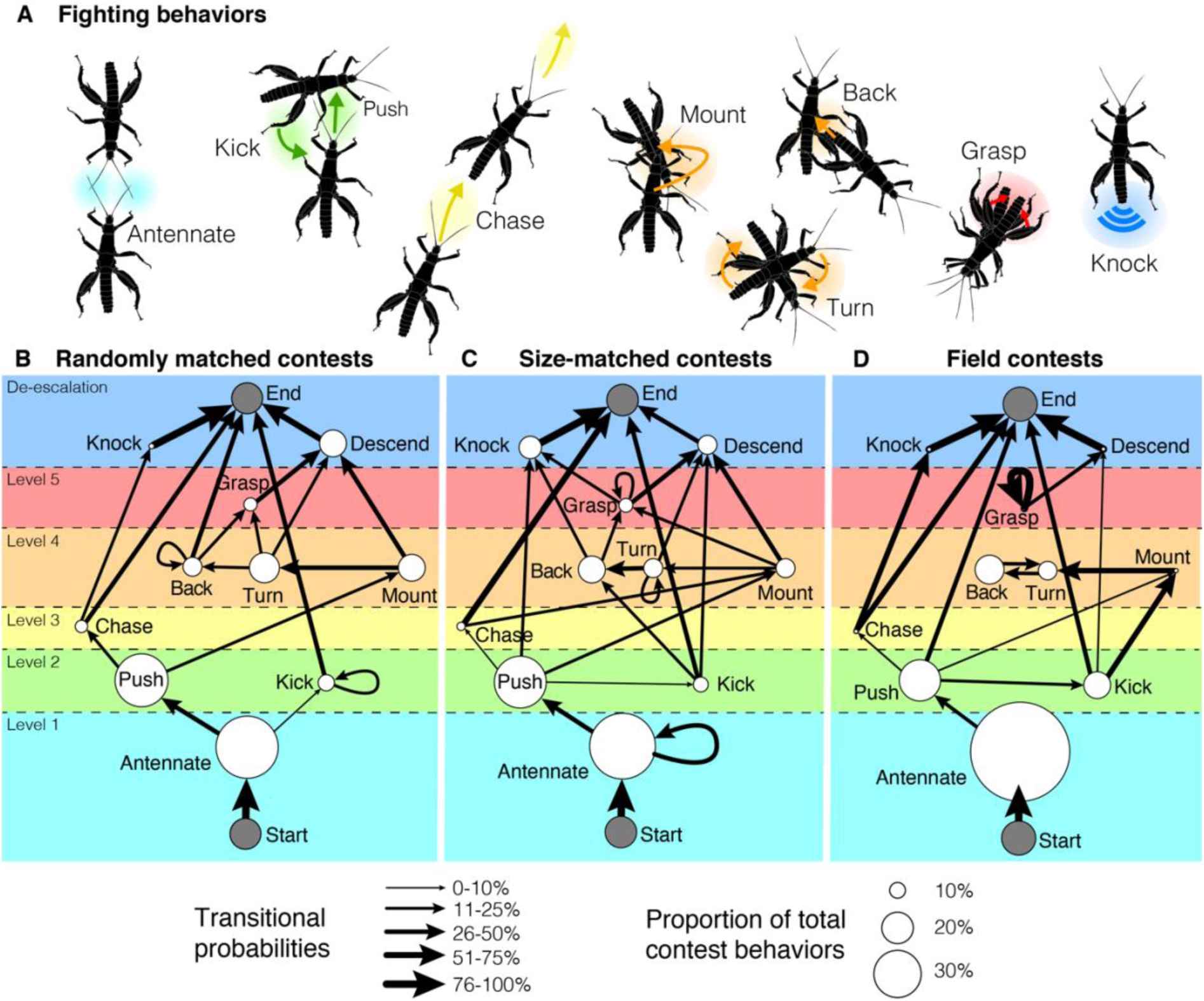
Contest behaviors (A) and sequential analysis in (B) randomly matched and (C) body size-matched lab contests, and (D) field contests. Contest behaviors progressed in phases (represented by different colors) of increasing intensity and escalation level (level 1 to 5) eventually leading to the resolution of the contest and its termination (de-escalation). Only significant transitions (i.e., occurring more often than expected randomly) are shown (arrows). The thickness of the arrows is proportional to transitional probability. No significant transition towards past phases that already occurred were observed. Individual behaviors are represented by circles, the size of which is scaled to the proportion of total contest behaviors represented by the given behavior.

More escalated behaviors (e.g., grasping) occurred more often in lab than field contests, likely because confined arenas limited retreat opportunities. Field fights typically ended after antennal contact or brief pushes (e.g., Video S3 in Boisseau et al. 2020), though the overall transition structure was similar across environments. As predicted by mutual assessment, size-matched contests showed a higher proportion of backward movements and grasping than randomly matched ones (8.0% vs. 6.4% and 4.4% vs. 4.0% of behaviors, respectively; Fig. 4B–C).

During grasping, males sat atop opponents and squeezed their hind femora—consistent with puncture injuries observed in both wild and lab individuals (Boisseau et al. 2020). Fighting behaviors suggest the use of tactile, chemical (antennal), and vibrational (abdominal thumping) cues for assessment. Hindlegs were never displayed or waved, indicating a function as combat tools rather than signaling structures.

### Correlational tests of assessment models

Contest costs decreased with mesothorax length asymmetry (χ² = 23.3, df = 1, p < 0.0001; Fig. 5A). Although predicted by mutual assessment, this pattern can also arise under self-assessment due to correlations between loser RHP and RHP asymmetry (Taylor and Elwood 2003). In randomly matched contests, contest costs increased with loser RHP (χ² = 6.87, df = 1, p = 0.009) but decreased with winner RHP (χ² = 4.85, df = 1, p = 0.028; Fig. 5B), ruling out pure self-assessment.

**Figure 5:**
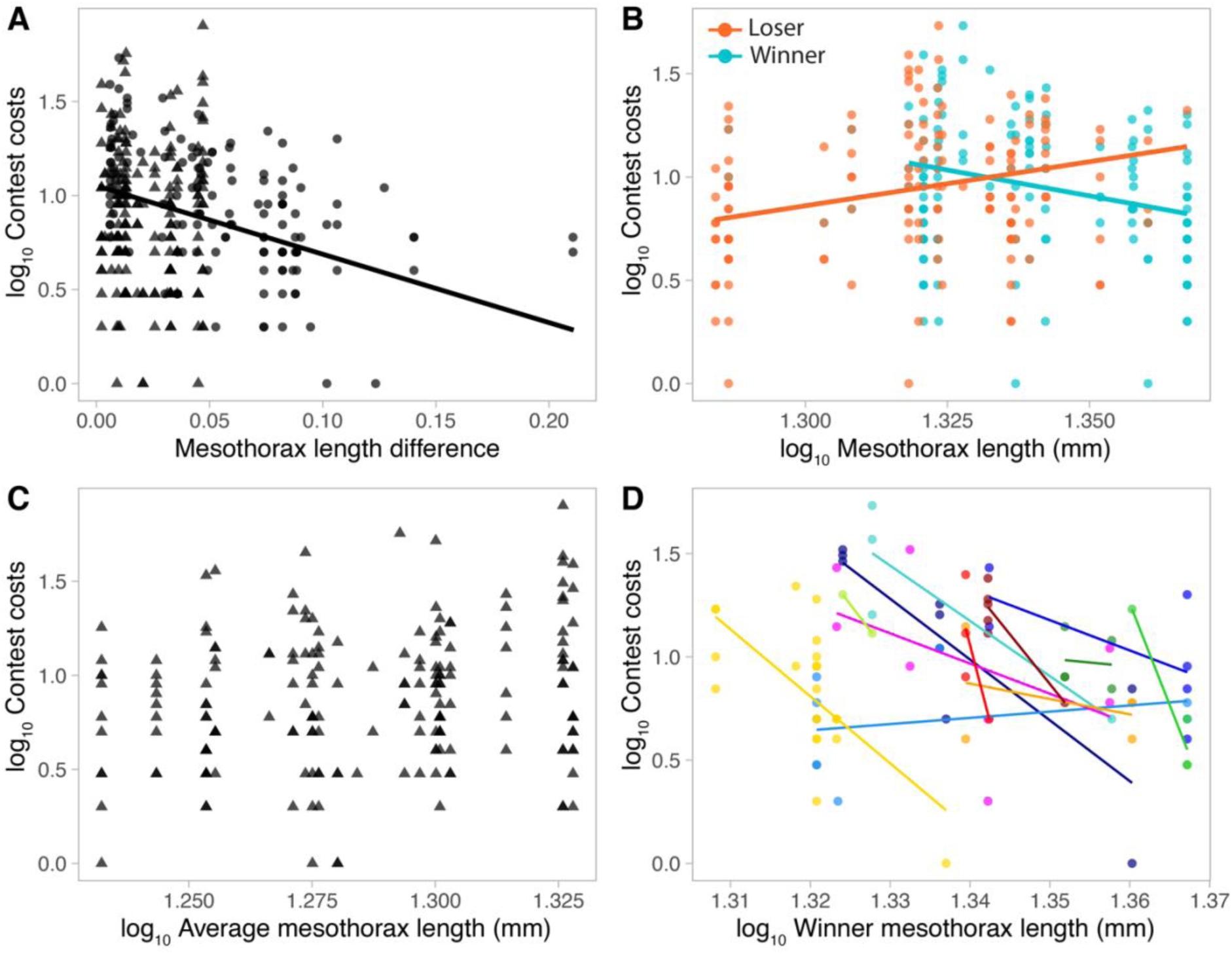
Correlational tests support mutual assessment during male-male contests. Relationships between contest costs and (A) mesothorax length asymmetry (i.e., absolute value of mesothorax length asymmetry index) in both randomly (circles) and size-matched contests (triangles), log_10_-corrected (B) loser (orange) and winner (blue) mesothorax length in randomly matched contests, (C) average contestant mesothorax length in size-matched contests. The repeated-testing approach is also presented for randomly matched contests (D): colors correspond to different focal losers which fought against several different contestants. Winner mesothorax length negatively affected contest costs for most focal losers.

In size-matched contests, average contestant RHP was only marginally correlated with contest costs (χ^2^ = 3.61, df = 1, p = 0.06; Fig. 5C), supporting mutual over cumulative assessment. Analyses of repeated contests for focal losers showed that contest costs decreased with winner RHP (χ^2^ = 8.40, df = 1, p = 0.004; Fig. 5D), and that individuals varied in both intercept (χ^2^ = 25.1, df = 1, p = 0.009) and slope (χ^2^ = 37.9, df = 1, p < 0.0001). For 11 of 12 focal losers, negative slopes indicated that males consistently employed mutual assessment (Fig. 5D).

### Sexual conflict and male-female interactions

Before copulation, males contacted females with their antennae, quickly mounted, aligned, and slid to the female’s right side, using the left hind leg to hold the abdomen (Fig. 6A). The male then passed the lower abdomen under the female to reach her genitalia. Females typically remained immobile, and no active resistance was observed. The male’s hind leg appeared to assist in bending the female’s abdomen to facilitate genital contact.

**Figure 6:**
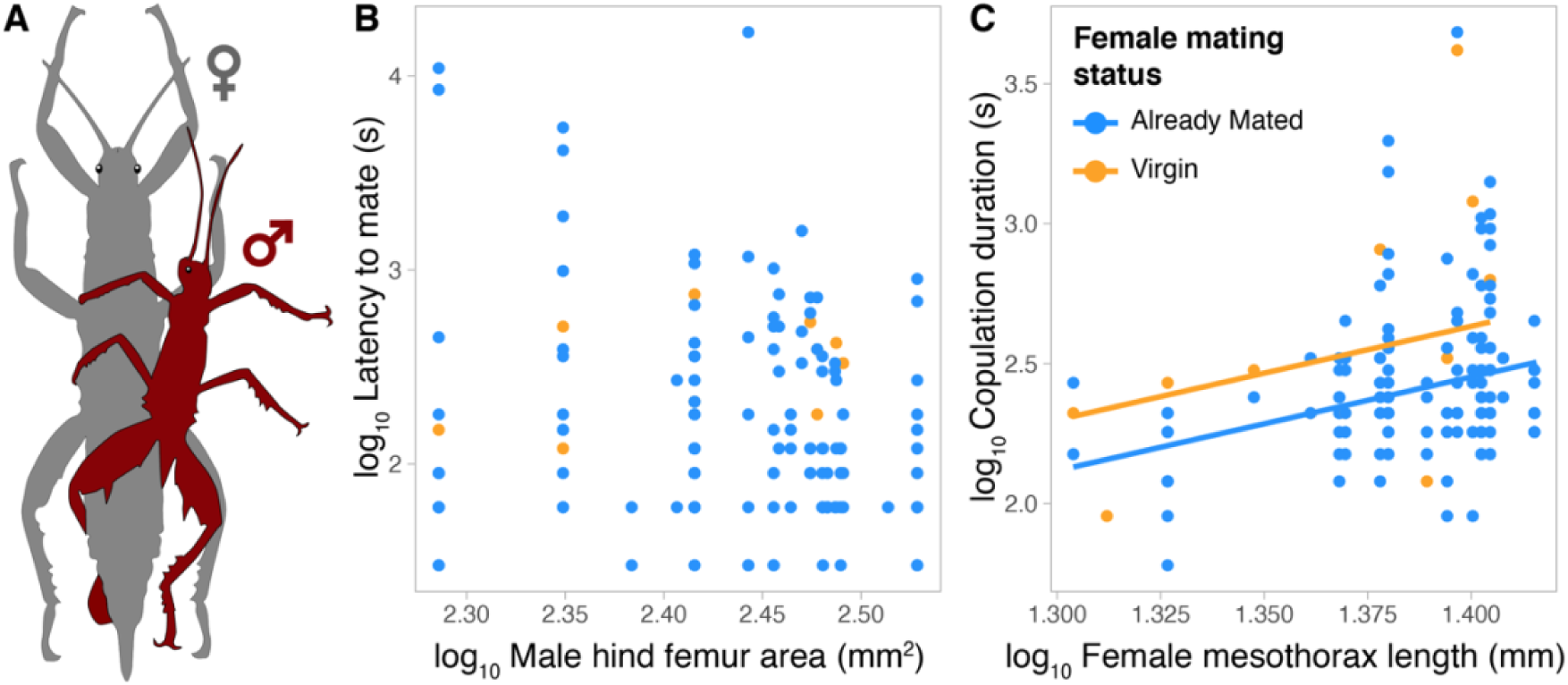
Male-female interactions in *E. calcarata*. (A) Typical copulation position. (B) Larger males with larger hind legs did not initiate copulation faster. (C) Males copulated for longer with both larger and virgin females.

Latency to mate was unaffected by male mesothorax length (χ² = 2.80, df = 1, p = 0.09), hind femur area (χ² = 1.85, df = 1, p = 0.17), female mesothorax length (χ² = 0.10, df = 1, p = 0.75), or female reproductive status (χ² = 1.63, df = 1, p = 0.20; Fig. 6B). Copulation duration was independent of male size (mesothorax χ² = 0.56, df = 1, p = 0.46; hind femur χ² = 0.01, df = 1, p = 0.92), but increased with female mesothorax length (χ² = 9.41, df = 1, p = 0.002) and was longer with virgin females (χ² = 6.32, df = 1, p = 0.01; Fig. 6C).

### Effect of mating on female fitness

Female size did not significantly affect egg number (F_1,25_= 0.79, p = 0.38) or average egg mass (F₁,₂₅ = 2.83, p = 0.10), but mated females laid more eggs than parthenogenetic ones (F_1,25_= 6.72, p = 0.02; Fig. 7A). Egg development was shorter in mated females (F_1,23_= 6.43, p = 0.018) and unaffected by female size (F_1,23_ = 0.23, p = 0.64; Fig. 7B). Egg-laying rate and hatching rate were independent of female size (F_1,25_ = 1.81, p = 0.19; F_1,23_ = 0.04, p = 0.84) and mating status (F_1,25_ = 0.23, p = 0.63; F_1,23_ = 2.73, p = 0.11; Fig. 7B–C).

**Figure 7:**
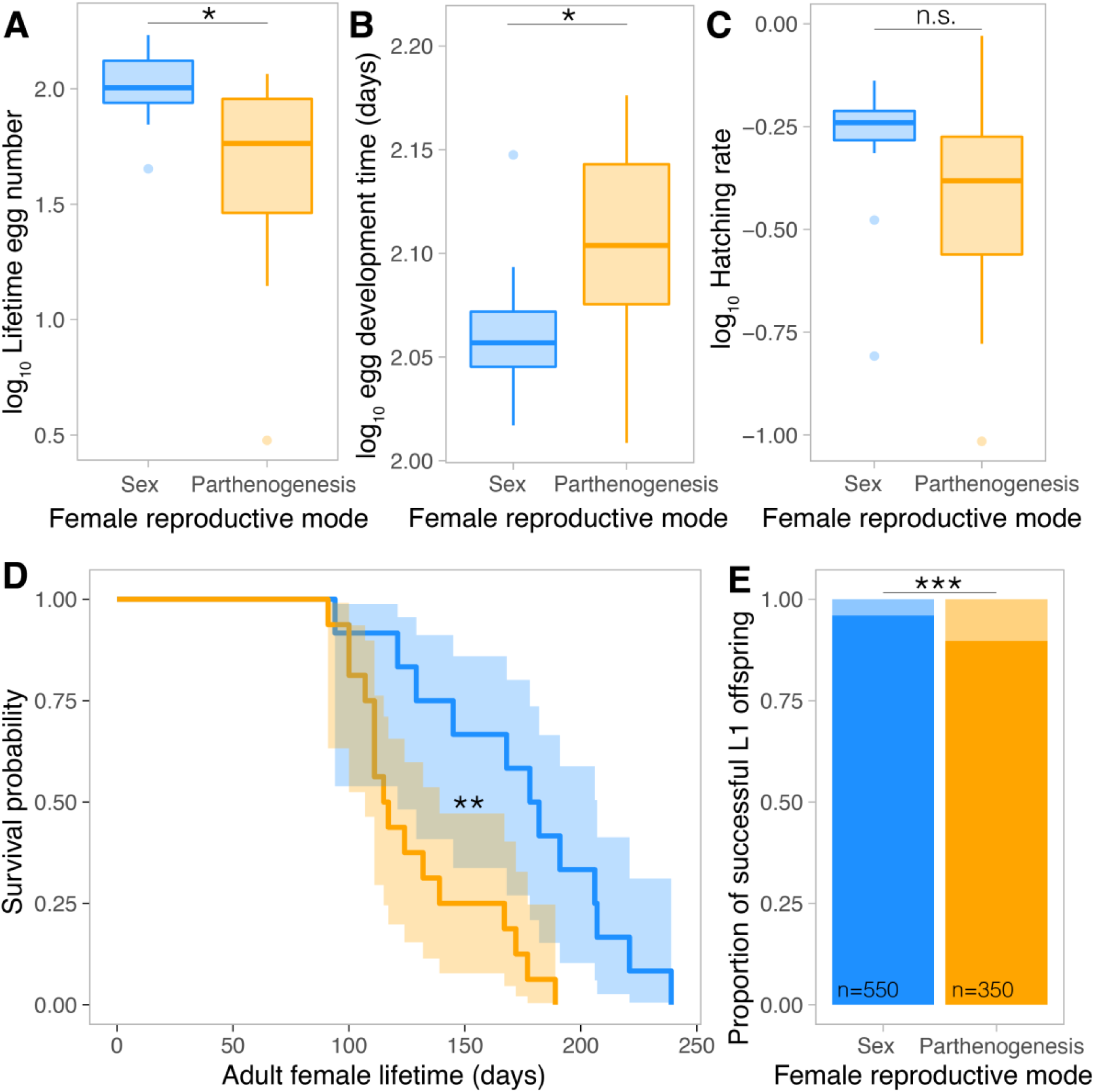
Effect of sex (blue) versus parthenogenesis (orange) on female fitness. Differences between mated and parthenogenetic females in terms of total lifetime egg number (**A**), egg development time (**B**), hatching rate (**C**), survival (**D**) and offspring survival during the first instar (**E**). Asterisks indicate significant differences (see Results for details).

Parthenogenetic females had shorter lifespans than mated females (116 vs. 180 days; χ^2^ = 9.7, df = 1, p = 0.002; Fig. 7D), and their offspring experienced higher first-instar mortality (10.2% vs. 4%; Fisher’s exact test: p = 0.0002; Fig. 7E).

## 4. Discussion

Our results suggest that the enlarged hindlegs and spines of *E. calcarata* males function primarily as cost-delivering tools of male-male combat, and not as agonistic deterrent (i.e., threat) signals or as tools to coerce females. During contests, larger males with longer and wider hindlegs were more likely to win and subsequently mate. Rival males did appear to mutually assess each other’s RHP during contests, likely using chemical, mechanical, and/or vibrational cues such as antennal contacts and abdominal thumping, but we found no evidence that hindlegs were used as visual or tactile signals. Hindlegs were used exclusively to deliver powerful squeezes to opponents, yet such instances were relatively rare because fights followed stereotyped escalating phases that often de-escalated before reaching the grasping stage.

The hindlegs of *E. calcarata* are secondary sexual traits that become strongly dimorphic in adults. Males use their hindlegs, equipped with sharp spines, to grab, squeeze, and puncture rivals’ body parts, primarily the hind femora, consistent with puncture wounds observed in the lab and field (Boisseau et al. 2020). These powerful squeezes are usually delivered while sitting atop the rival (Fig. 4A). Although some insects with enlarged hindlegs adopt a backwards-facing fighting style (Procter et al. 2012; O’Brien et al. 2017a; Fea and Holwell 2018; Rink et al. 2019), such behavior in *E. calcarata* and *E. horrida* was extremely rare (<1% of contests) (Clail 1988). Many weapon systems, like leaf-footed bugs (O’Brien and Boisseau 2018) and frog-legged beetles (O’Brien et al. 2017b), show hyperallometric scaling of hindlegs. Hyperallometry typically evolves when weapons also signal RHP, since steep allometry slopes exaggerate male differences making them easier for receivers to assess (Eberhard et al. 2018; O’Brien et al. 2018; McCullough and O’Brien 2022). However, hyperallometry can weaken the weapons of the largest individuals if the out-lever arm grows disproportionately relative to the in-lever arm (“paradox of the weakening combatant”) (Levinton and Allen 2005; O’Brien and Boisseau 2018). Isometry, on the other hand, typically preserves mechanical advantage by keeping lever components proportional (O’Brien and Boisseau 2018). The enlarged hindlegs of *E. calcarata*, like the hindleg weapons of cave wētās (Fea and Holwell 2018) and monkey beetles (Rink et al. 2019), scale proportionately with body size (Bonduriansky 2007), consistent with a function as a pure combat tool. The relatively large, costly femoral flexor muscles of *E. calcarata* further highlight the importance of squeezing strength (O’Brien et al. 2019).

Body size was the main predictor of contest outcome and used as a proxy for RHP. After accounting for body size, weapon size and body condition had no significant effect on winning in either randomly- or size-matched contests, likely because weapon and body size are tightly correlated (Painting and Holwell 2014; del Sol et al. 2020). In body size-matched contests, only residency affected outcomes, with resident males more likely to win, and the effect of weapon size remained non-significant. This suggests disproportionately large hindlegs do not confer a combat advantage, possibly due to higher maintenance costs (O’Brien et al. 2019) or reduced mechanical performance (O’Brien and Boisseau 2018). This contrasts with systems where weapon size outweighs body size in predicting fighting success (Bridge et al. 2000; Lailvaux et al. 2005; Fea and Holwell 2018). In those systems, other weapon traits like reach may be more critical than strength, which depends on musculature, stamina, and lever mechanics.

Although weapon size was not a stronger predictor of RHP than body size, this does not exclude a signaling role during combat, which would require opponents to assess each other’s RHP—consistent with opponent-only or mutual assessment models. Fights escalated less when size disparities were large, more when losers were larger, and less when winners were larger, ruling out pure self-assessment (Arnott and Elwood 2009; Green and Patek 2018; Fig. 1B). Contest costs and escalation levels were similar between large and small size-matched males, supporting mutual rather than cumulative assessment (Pinto et al. 2019; Fig. 1C). Repeated contests with small focal losers showed greater escalation when winners were smaller, again indicating widespread mutual assessment (Chapin et al. 2019; Fig. 1D). Behavioral sequence analyses confirmed that contests escalated through distinct phases, with size-matched males more often reaching the most intense stages (Green and Patek 2018; Fig. 1E). Together, our results indicate that *E. calcarata* males assess rival size and RHP during combat and use this information to decide whether to persist or retreat.

Males appeared to assess rivals primarily through antennal contact, which consistently initiated contests, suggesting the use of mechanical or chemical cues to gauge opponent size early on. The specific cues remain unknown (Arnott and Elwood 2009), but cuticular hydrocarbons are good candidates as they have been shown to signal body size or dominance in other arthropods (Thomas and Simmons 2011; Steiger et al. 2013; Lane et al. 2016). These contacts were not directed toward the hindlegs, and no behaviors such as leg waving or backward displays indicated a signaling role for the hindlegs, unlike in other weapon systems (Clutton-Brock et al. 1979; Miyatake 1993; Jennions 1996; Katsuki et al. 2014; McCullough et al. 2016). Winners frequently thumped their abdomen on the substrate after contests—a behavior also reported in other phasmids (James 1981; Delfosse 2003)—which may transmit information about body size or RHP via vibrational substrate cues (De Luca and Morris 1998; Cocroft and Rodríguez 2005). Because thumping occurred almost exclusively in winners, it likely represents a victory display (Bower 2005; Kelly 2006; Chen et al. 2014), possibly advertising dominance to nearby males or discouraging re-engagement by defeated rivals (Mesterton-Gibbons and Sherratt 2006; Chen et al. 2017). Given the high male densities on tree trunks (Boisseau et al. 2020), such signals could reach multiple bystanders and influence future contest dynamics.

Exaggerated male hindlegs are often used to coerce females in insects (Rowe et al. 2006; Haley and Gray 2012; Burrows 2020). In *E. calcarata*, males use their hindlegs to grasp females and position their abdomen during copulation (Hsiung 1987), suggesting a potential role in overcoming resistance. However, mating latency was unaffected by male or female size or by female mating status, and females showed no behavioral resistance—typically freezing upon contact and remaining immobile. We showed that avoiding mating and reproducing via parthenogenesis proved costly: unmated females had shorter adult lifespans, laid fewer eggs with longer development times, and produced offspring with higher mortality (Burke et al. 2015). These costs are consistent with virgin females not resisting males more than mated females. In the field, females mated with several males nightly without resisting (Boisseau et al. 2020), suggesting that multiple mating may offer benefits or that resisting repeated copulation attempts is more costly than mating (“convenience polyandry”; Rowe 1992; Cordero and Andrés 2002; Arnqvist and Rowe 2005). Given the high densities of males on tree trunks, tolerance of copulation may reduce harassment costs. Nonetheless, under lower male densities, resistance could occur (Rowe 1992), so a coercive function of the male hindlegs cannot be entirely ruled out.

## 5. Conclusion

Exaggerated male weapons are rare in Phasmatodea but have evolved convergently in at least three lineages (Buckley et al. 2009; Boisseau et al. 2020; Emberts and Wiens 2021). Building on previous field work linking this evolution to communal roosting and defense-based polygyny (Boisseau et al. 2020), our study shows that larger males with proportionately larger hindlegs achieve greater fighting and mating success. In parallel, we found no evidence that hindlegs function coercively during copulation, despite their use in grasping females, indicating that exaggeration is driven mainly by male-male competition. These hindlegs serve as cost-delivering weapons rather than threat signals, consistent with their isometric scaling with body size (Eberhard et al. 2018; O’Brien et al. 2018; McCullough and O’Brien 2022). Although males appear to mutually assess opponents’ size and fighting ability, this likely relies on non-visual cues, independent of the hindlegs. By integrating field observations and analyses of contest outcomes, costs, behaviors, and mating interactions, we were able to characterize the function of this sexually dimorphic trait and shed light on its evolution and scaling pattern.

## 6. Author contributions

R.P.B and D.J.E conceived of the study. R.P.B conducted the experiments in the field and in the lab, reared the study animals, performed the statistical analyses and wrote the initial version of the manuscript. R.P.B and D.J.E contributed to editing and revising subsequent versions of the manuscript.

## 7. Acknowledgements

We thank M. Ero, L. Bonneau, S. Makai, R. Dikrey, B. Sapau, P. Mana, S. Komda, G. Gumbira, R. Uker, T. Manjobie and T. Batari for assistance in the field in Papua New Guinea; C. Thomas-Bulle for helping with the experiments in the lab and for helpful discussions and feedback on the manuscript; the Missoula Butterfly house and Insectarium for providing the initial captive population of *E. calcarata*; P. Green for kindly providing R scripts to run the behavioral sequence analyses; C. Allen for help with insect rearing and logistics.

## 8. Data availability

All data and code will be made publicly available on Zenodo upon acceptance.

## 9. Conflicts of interest

The authors declare no competing interests.

## 10. Funding

The field part of the project was funded by an early career grant from the National Geographic Society to R.P.B (WW-255ER-17) and by the Papua New Guinea Oil Palm Research Association (PNGOPRA). The lab part of the project was funded by the NSF IOS–1456133 & IOS-2015907 (D.J.E).

## Notes

### Competing Interest Statement

The authors have declared no competing interest.

## References

Andersson, M. 1994. Sexual Selection. Princeton University Press, Princeton, NJ, USA.

Arnott, G., and R. W. Elwood. 2009. Assessment of fighting ability in animal contests. Animal Behaviour 77:991–1004.

Arnott, G., and R. W. Elwood. 2008. Information gathering and decision making about resource value in animal contests. Animal Behaviour 76:529–542.

Arnqvist, G., and L. Rowe. 2005. Sexual conflict. Princeton University Press, Princeton, New Jersey, USA.

Bedford, G. O. 1976. Defensive behaviour of the New Guinea stick insect *Eurycantha* (Phasmatodea: Phasmatidae: Eurycanthinae). Proceedings of the Linnean Society of New South Wales 100:218–222.

Berglund, A., A. Bisazza, and A. Pilastro. 1996. Armaments and ornaments: an evolutionary explanation of traits of dual utility. Biological Journal of the Linnean Society 58:385–399.

Bildstein, K. L., S. G. McDowell, and I. L. Brisbin. 1989. Consequences of sexual dimorphism in sand fiddler crabs, *Uca pugilator*: differential vulnerability to avian predation. Animal Behaviour 37:133–139.

Boisseau, R. P., M. M. Ero, S. Makai, L. J. G. Bonneau, and D. J. Emlen. 2020. Sexual dimorphism divergence between sister species is associated with a switch in habitat use and mating system in thorny devil stick insects. Behavioural Processes 181:104263.

Bonduriansky, R. 2007. Sexual selection and allometry: A critical reappraisal of the evidence and ideas. Evolution 61:838–849.

Bower, J. L. 2005. The occurrence and function of victory displays within communication networks. P. 115—126 in P. McGregor, ed. Animal Communication Networks. Cambridge University press, Cambridge, UK.

Bridge, A. P., R. W. Elwood, and J. T. A. Dick. 2000. Imperfect assessment and limited information preclude optimal strategies in male-male fights in the orb-weaving spider *Metellina mengei*. Proceedings of the Royal Society B 267:273–279.

Briffa, M., I. C. W. Hardy, M. P. Gammell, D. J. Jennings, D. D. Clarke, and M. Goubault. 2013. Analysis of animal contest data. Pp. 47–85 in I. C. W. Hardy and M. Briffa, eds. Animal Contests. Cambridge University press, Cambridge, United Kingdom.

Buckley, T. R., D. Attanayake, and S. Bradler. 2009. Extreme convergence in stick insect evolution: phylogenetic placement of the Lord Howe Island tree lobster. Proceedings of the Royal Society B: Biological Sciences 276:1055–1062.

Burke, N. W., and R. Bonduriansky. 2022. Sexually but not parthenogenetically produced females benefit from mating in a stick insect. Functional Ecology, doi: 10.1111/1365-2435.14095.

Burke, N. W., A. J. Crean, and R. Bonduriansky. 2015. The role of sexual conflict in the evolution of facultative parthenogenesis: A study on the spiny leaf stick insect. Animal Behaviour 101:117–127.

Burrows, M. 2020. Do the enlarged hind legs of male thick-legged flower beetles contribute to take-off or mating? Journal of Experimental Biology 223:jeb212670.

Carlberg, U. 1989. Aspects of Defensive Behaviour of *Eurycantha calcarata* Lucas Females and the Evolution of Scorpion Mimicry in the Phasmida (Insecta). Biologisches Zentralblatt 108:257–262.

Chapin, K. J., P. E. C. Peixoto, and M. Briffa. 2019. Further mismeasures of animal contests: A new framework for assessment strategies. Behavioral Ecology 30:1177–1185.

Chen, P. Z., L. R. Carrasco, and P. K. L. Ng. 2014. Post-contest stridulation used exclusively as a victory display in mangrove crabs. Ethology 120:532–539. John Wiley & Sons, Ltd.

Chen, P. Z., R. L. Carrasco, and P. K. L. Ng. 2017. Mangrove crab uses victory display to “browbeat” losers from re-initiating a new fight. Ethology 123:981–988. John Wiley & Sons, Ltd.

Clail, I. 1988. A behavioural study of Eurycantha calcarata. Stirling University.

Clutton-Brock, T. H., S. D. Albon, R. M. Gibson, and F. E. Guinness. 1979. The logical stag: adaptive aspects of fighting in red deer (*Cervus elaphus* L.). Animal Behaviour 27:211–225. Academic Press.

Clutton-Brock, T., and G. A. Parker. 1995. Sexual coercion in animal societies. Animal Behaviour 49:1345–1365.

Cocroft, R. B., and R. L. Rodríguez. 2005. The Behavioral Ecology of Insect Vibrational Communication. BioScience 55:323–334. Narnia.

Cordero, A., and J. A. Andrés. 2002. Male coercion and convenience polyandry in a calopterygid damselfly. Journal of Insect Science 2:1–7.

Csardi, G., and T. Nepusz. 2006. The igraph software package for complex network research. InterJournal, complex systems 1695:1–9.

De Luca, P. A., and G. K. Morris. 1998. Courtship communication in meadow katydids: female preference for large male vibrations. Behaviour 135:777–794.

del Sol, J. F., Y. Hongo, R. Boisseau, G. Berman, C. E. Allen, and Douglas. J. Emlen. 2020. Population differences in the strength of sexual selection match relative weapon size in the Japanese rhinoceros beetle, *Trypoxylus dichotomus* (Coleoptera: Scarabaeidae). Evolution 75:394–413.

Delfosse, E. 2003. Taxonomie, répartition, élevage, émission et réception de sons chez le Phasme épineux « marteleur » : *Aretaon* (*Aretaon*) *asperrimus* (Redtenbacher, 1906) (Insecta Orthopteroidea Phasmatodea Areolatae Bacillidae Heteropteryginae Obrimini. Bulletin de Phyllie 16:16–30.

Eberhard, W. G., S. José, C. Rica, R. Lucas Rodríguez, B. A. H. Alexander, B. Speck, H. Miller, R. L. Rodríguez, B. A. Huber, B. Speck, H. Miller, B. A. Buzatto, and G. Machado. 2018. Sexual Selection and Static Allometry: the Importance of Function. The Quarterly Review of Biology 93:207–250.

Elwood, R. W., and G. Arnott. 2013. Assessments in contests are frequently assumed to be complex when simple explanations will suffice. Animal Behaviour 86:e8–e12.

Elwood, R. W., and G. Arnott. 2012. Understanding how animals fight with Lloyd Morgan’s canon. Animal Behaviour 84:1095–1102.

Emberts, Z., and J. J. Wiens. 2021. Do sexually selected weapons drive diversification? Evolution 75:2411–2424.

Emlen, D. J. 2008. The Evolution of Animal Weapons. Annual Review of Ecology, Evolution, and Systematics 39:387–413.

Enquist, M., and O. Leimar. 1983. Evolution of Fighting Behaviour : Decision Rules and Assessment of Relative Strength. Journal of Theoretical Biology 102:387–410.

Enquist, M., O. Leimar, T. Ljungberg, Y. Mallner&, and N. Segerdahl. 1990. A test of the sequential assessment game: fighting in the cichlid fish *Nannacara anomala*. Animal Behaviour 40:1–14.

Fea, M., and G. Holwell. 2018. Combat in a cave-dwelling wētā (Orthoptera: Rhaphidophoridae) with exaggerated weaponry. Animal Behaviour 138:85–92.

Goubault, M., and M. Decuignière. 2012. Previous Experience and Contest Outcome: Winner Effects Persist in Absence of Evident Loser Effects in a Parasitoid Wasp. The American Naturalist 180:364–371.

Green, P. A., and S. N. Patek. 2018. Mutual assessment during ritualized fighting in mantis shrimp (Stomatopoda). Proceedings of the Royal Society B: Biological Sciences 285:20172542.

Haley, E. L., and D. A. Gray. 2012. Mating behavior and dual-purpose armaments in a camel cricket. Ethology 118:49–56.

Hardy, I. C. W., and M. Briffa. 2013. Animal Contests. Cambridge University press, Cambridge, United Kingdom.

Hsiung, C.-C. 1987. Aspects of the biology of the Melanesian stick-insect *Eurycantha calcarata* Lucas (Cheleutoptera: Phasmatidae). Journal of Natural History 21:1241–1258.

Hsu, Y., R. L. Earley, and L. L. Wolf. 2006. Modulation of aggressive behaviour by fighting experience: mechanisms and contest outcomes. Biological Reviews 81:33–74.

James, T. 1981. *Anisomorpha buprestoides* noises. The Phasmid Study Group Newsletter 6:2.

Jennions, M. D. 1996. Residency and size affect fight duration and outcome in the fiddler crab *Uca annulipes*. Biological Journal of the Linnean Society 57:293–306. Oxford University Press (OUP).

Katsuki, M., T. Yokoi, K. Funakoshi, and N. Oota. 2014. Enlarged hind legs and sexual behavior with male-male interaction in *Sagra femorata* (Coleoptera : Chrysomelidae). Entomological news 124:211–220.

Kelly, C. D. 2006. Fighting for harems : assessment strategies during male - male contests in the sexually dimorphic Wellington tree weta. Animal behaviour 72:727–736.

Koczur, W., J. Szwedo, and M. Gołębiowski. 2024. The defensive secretion of Eurycantha calcarata (Phasmida: Lonchodidae) - chemical composition and method of collection. EJE 121:360–368. EJE.

Lailvaux, S. P., J. Hathway, J. Pomfret, and R. J. Knell. 2005. Horn size predicts physical performance in the beetle *Euoniticellus intermedius* (Coleoptera: Scarabaeidae). Functional Ecology 19:632–639.

Lane, S. M., A. W. Dickinson, T. Tregenza, and C. M. House. 2016. Sexual Selection on male cuticular hydrocarbons via male–male competition and female choice. Journal of Evolutionary Biology 29:1346–1355. John Wiley & Sons, Ltd.

Lane, S. M., and E. L. McCullough. 2025. The prevalence of weapon damage: a proportional meta-analysis. Animal Behaviour 222:123117.

Levinton, J. S., and B. J. Allen. 2005. The paradox of the weakening combatant: Trade-off between closing force and gripping speed in a sexually selected combat structure. Functional Ecology 19:159–165.

Lovich, J. E., and J. W. Gibbons. 1992. A review of techniques for quantifying sexual size dimorphism. Growth, Development and Aging 56:269–281.

Maynard Smith, J. 1982. Evolution and the Theory of Games. Cambridge University press, Cambridge, United Kingdom.

Maynard Smith, J. 1974. The theory of games and the evolution of animal conflicts. Journal of Theoretical Biology 47:209–221.

Maynard Smith, J., and G. A. Parker. 1976. The logic of asymmetric contests. Animal Behaviour 24:159–175.

Maynard Smith, J., and G. R. Price. 1973. Logic of animal conflict. Nature 246:15–18.

McCullough, E. L., C. W. Miller, and D. J. Emlen. 2016. Why sexually selected weapons are not ornaments. Trends in Ecology and Evolution 31:742–751.

McCullough, E. L., and D. M. O’Brien. 2022. Variation in allometry along the weapon-signal continuum. Evolutionary Ecology 36:591–604.

McLain, D. K., A. E. Pratt, and A. S. Berry. 2003. Predation by red-jointed fiddler crabs on congeners: interaction between body size and positive allometry of the sexually selected claw. Behavioral Ecology 14:741–747.

Mesterton-Gibbons, M., J. H. Marden, and L. A. Dugatkin. 1996. On wars of attrition without assessment. Journal of Theoretical Biology 181:65–83.

Mesterton-Gibbons, M., and T. N. Sherratt. 2006. Victory displays: a game-theoretic analysis. Behavioral Ecology 17:597–605. Oxford Academic.

Miyatake, T. 1993. Male-male aggressive behavior is changed by body size difference in the leaf-footed plant bug, *Leptoglossus australis*, Fabricius (Heteroptera: Coreidae). Journal of Ethology 11:63–65.

Muramatsu, D. 2011. The function of the four types of waving display in *Uca lactea*: effects of audience, sand structure, and body size. Ethology 117:408–415.

O’Brien, D. M., C. E. Allen, M. J. Van Kleeck, D. Hone, R. Knell, A. Knapp, S. Christiansen, and D. J. Emlen. 2018. On the evolution of extreme structures: static scaling and the function of sexually selected signals. Animal Behaviour 144:95–108.

O’Brien, D. M., and R. P. Boisseau. 2018. Overcoming mechanical adversity in extreme hindleg weapons. Plos One 13:e0206997.

O’Brien, D. M., R. P. Boisseau, M. Duell, E. McCullough, E. C. Powell, U. Somjee, S. Solie, A. J. Hickey, G. I. Holwell, C. J. Painting, and D. J. Emlen. 2019. Muscle mass drives cost in sexually selected arthropod weapons. Proceedings of the Royal Society B 286:20191063. The Royal Society.

O’Brien, D. M., M. Katsuki, and D. J. Emlen. 2017a. Selection on an extreme weapon in the frog-legged leaf beetle (*Sagra femorata*). Evolution 71:2584–2598.

O’Brien, D. M., M. Katsuki, and D. J. Emlen. 2017b. Selection on an extreme weapon in the frog-legged leaf beetle (*Sagra femorata*). Evolution 71:2584–2598.

Painting, C. J., and G. I. Holwell. 2014. Exaggerated rostra as weapons and the competitive assessment strategy of male giraffe weevils. Behavioral Ecology 25:1223–1232. Oxford Academic.

Parker, G. A. 1974. Assessment strategy and the evolution of animal conflicts. Journal of Theoretical Biology 47:223–243.

Payne, R. J. H. 1998. Gradually escalating fights and displays : the cumulative assessment model. Animal Behaviour 56:651–662.

Payne, R. J. H., and M. Pagel. 1996. Escalation and time costs in displays of endurance. Journal of Theoretical Biology 183:185–193.

Pinheiro, J., D. Bates, S. DebRoy, D. Sarkar, and R Core Team. 2021. nlme: linear and nonlinear mixed effects models. R package version 3.1–162.

Pinto, N. S., A. V. Palaoro, and P. E. C. Peixoto. 2019. All by myselfMeta-analysis of animal contests shows stronger support for self than for mutual assessment models. Biological Reviews 55.

Pradhan, G. R., and C. P. Van Schaik. 2009. Why do females find ornaments attractive? The coercion-avoidance hypothesis. Biological Journal of the Linnean Society 96:372–382.

Pratt, A. E., D. Kelly McLain, and G. R. Lathrop. 2003. The assessment game in sand fiddler crab contests for breeding burrows. Animal Behaviour 65:945–955.

Procter, D. S., a. J. Moore, and C. W. Miller. 2012. The form of sexual selection arising from male-male competition depends on the presence of females in the social environment. Journal of Evolutionary Biology 25:803–812.

R Core Team. 2023. R: A Language and Environment for Statistical Computing. R Foundation for Statistical Computing, Vienna, Austria.

Rico-Guevara, A., and K. J. Hurme. 2019. Intrasexually selected weapons. Biological Reviews 94:60–101.

Rink, A. N., R. Altwegg, S. Edwards, R. C. K. Bowie, and J. F. Colville. 2019. Contest dynamics and assessment strategies in combatant monkey beetles (Scarabaeidae: Hopliini). Behavioral Ecology 30:713–723. Oxford Academic.

Rodríguez, R. L., and W. G. Eberhard. 2019. Why the static allometry of sexually-selected traits is so variable: the importance of function. Integrative and Comparative Biology 59:1290–1302.

Rometsch, S. J., J. Torres-Dowdall, G. Machado-Schiaffino, N. Karagic, and A. Meyer. 2021. Dual function and associated costs of a highly exaggerated trait in a cichlid fish. Ecology and Evolution 11:17496–17508.

Rowe, L. 1992. Convenience polyandry in a water strider: foraging conflicts and female control of copulation frequency and guarding duration. Animal Behaviour 44:189–202.

Rowe, L., K. P. Westlake, and D. C. Currie. 2006. Functional significance of elaborate secondary sexual traits and their evolution in the water strider genus *Rheumatobates*. Canadian Entomologist 138:568–577.

Rutte, C., M. Taborsky, and M. W. G. Brinkhof. 2006. What sets the odds of winning and losing? Trends in Ecology & Evolution 21:16–21.

Schneider, C. A., W. S. Rasband, and K. W. Eliceiri. 2012. NIH Image to ImageJ: 25 years of image analysis. Nature Methods 9:671–675.

Shuker, D. M., and L. W. Simmons. 2014. The evolution of insect mating systems. Oxford University Press, Oxford, United Kingdom.

Steiger, S., G. D. Ower, J. Stökl, C. Mitchell, J. Hunt, and S. K. Sakaluk. 2013. Sexual selection on cuticular hydrocarbons of male sagebrush crickets in the wild. Proceedings of the Royal Society B: Biological Sciences 280.

Taylor, P. W., and R. W. Elwood. 2003. The mismeasure of animal contests. Animal Behaviour 65:1195–1202.

Thomas, M. L., and L. W. Simmons. 2011. Short-term phenotypic plasticity in long-chain cuticular hydrocarbons. Proceedings of the Royal Society B: Biological Sciences 278:3123–3128.

