## Supplementary Information for "Exaggerated male hindlegs function as pure weapons of male–male combat in thorny devil stick insects"

**This file includes:**

- Tables S1
- Figures S1-S3
- Legends for Datasets S1

### Supplementary tables

**Table S1 :** Analyses of the scaling relationships between front femur length, hind femur length, hind femur width, hind femur area and mesothorax length (~body size) throughout development and across sexes. The table shows the results of best linear mixed-effects models (accounting for individual ID as a random effect) and associated type I ANOVA. Departure from isometry (i.e., slope= 1 for linear traits, slope= 2 for surface traits) was tested in adults using 95% confidence intervals (between brackets) around estimated regression slopes.

| Response variable | N | Explanatory variables | F | df1 | df2 | P | Isometric slope | Adult slope |
|---|---|---|---|---|---|---|---|---|
| $Log_{10}$(Front femur length) | 254 | $log_{10}$(Mesothorax length) | 18119 | 1 | 133 | <.0001 | 1 | 0.83 [0.70; 0.96] |
| | | instar | 119 | 3 | 133 | <.0001 | | |
| | | sex | 38 | 1 | 109 | <.0001 | | |
| | | instar:sex | 9 | 3 | 133 | <.0001 | | |
| | | $log_{10}$(Mesothorax length):instar | 4 | 3 | 133 | 0.01 | | |
| $Log_{10}$(Hind femur length) | 253 | $log_{10}$(Mesothorax length) | 25047 | 1 | 132 | <.0001 | 1 | 0.97 [0.82; 1.12] |
| | | instar | 345 | 3 | 132 | <.0001 | | |
| | | sex | 261 | 1 | 109 | <.0001 | | |
| | | instar:sex | 99 | 3 | 132 | <.0001 | | |
| | | $log_{10}$(Mesothorax length):instar | 8 | 3 | 132 | <.0001 | | |
| $Log_{10}$(Hind femur width) | 253 | $log_{10}$(Mesothorax length) | 7821 | 1 | 135 | <.0001 | 1 | 1.05 [0.81; 1.29] |
| | | instar | 167 | 3 | 135 | <.0001 | | |
| | | sex | 447 | 1 | 109 | <.0001 | | |
| | | instar:sex | 129 | 3 | 135 | <.0001 | | |
| $Log_{10}$(Hind femur area) | 253 | $log_{10}$(Mesothorax length) | 17763 | 1 | 132 | <.0001 | 2 | 2.02 [1.69; 2.36] |
| | | instar | 300 | 3 | 132 | <.0001 | | |
| | | sex | 526 | 1 | 109 | <.0001 | | |
| | | instar:sex | 164 | 3 | 132 | <.0001 | | |
| | | $log_{10}$(Mesothorax length):instar | 5 | 3 | 132 | 0.002 | | |

#### Supplementary figures

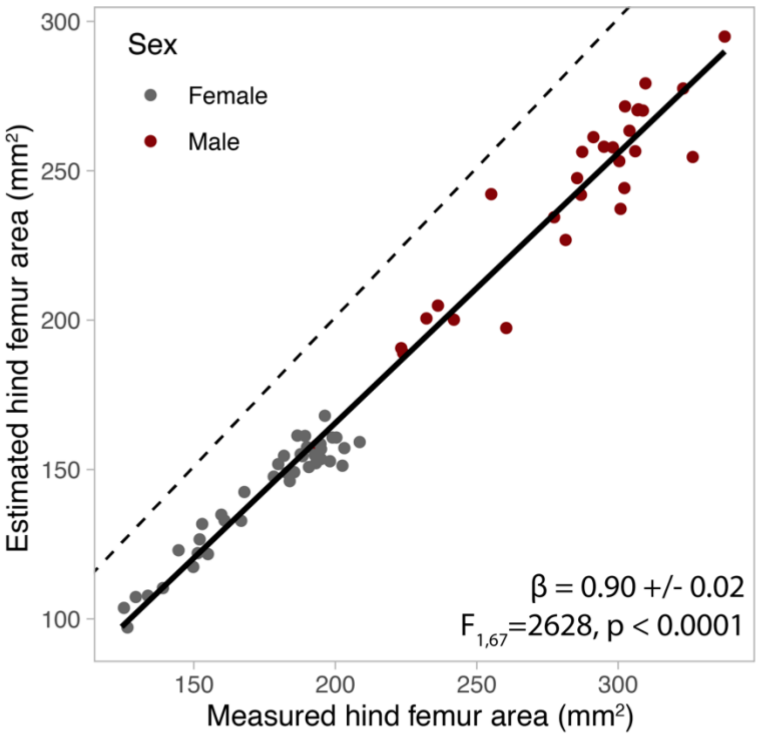


**Figure S1:** Correlation between estimated (i.e., the area of the ellipse of major diameter hind femur length and minor diameter hind femur width) and measured hind femur area (i.e., the area obtained by drawing the lateral outline of the femur). The dashed line represents the equality line.

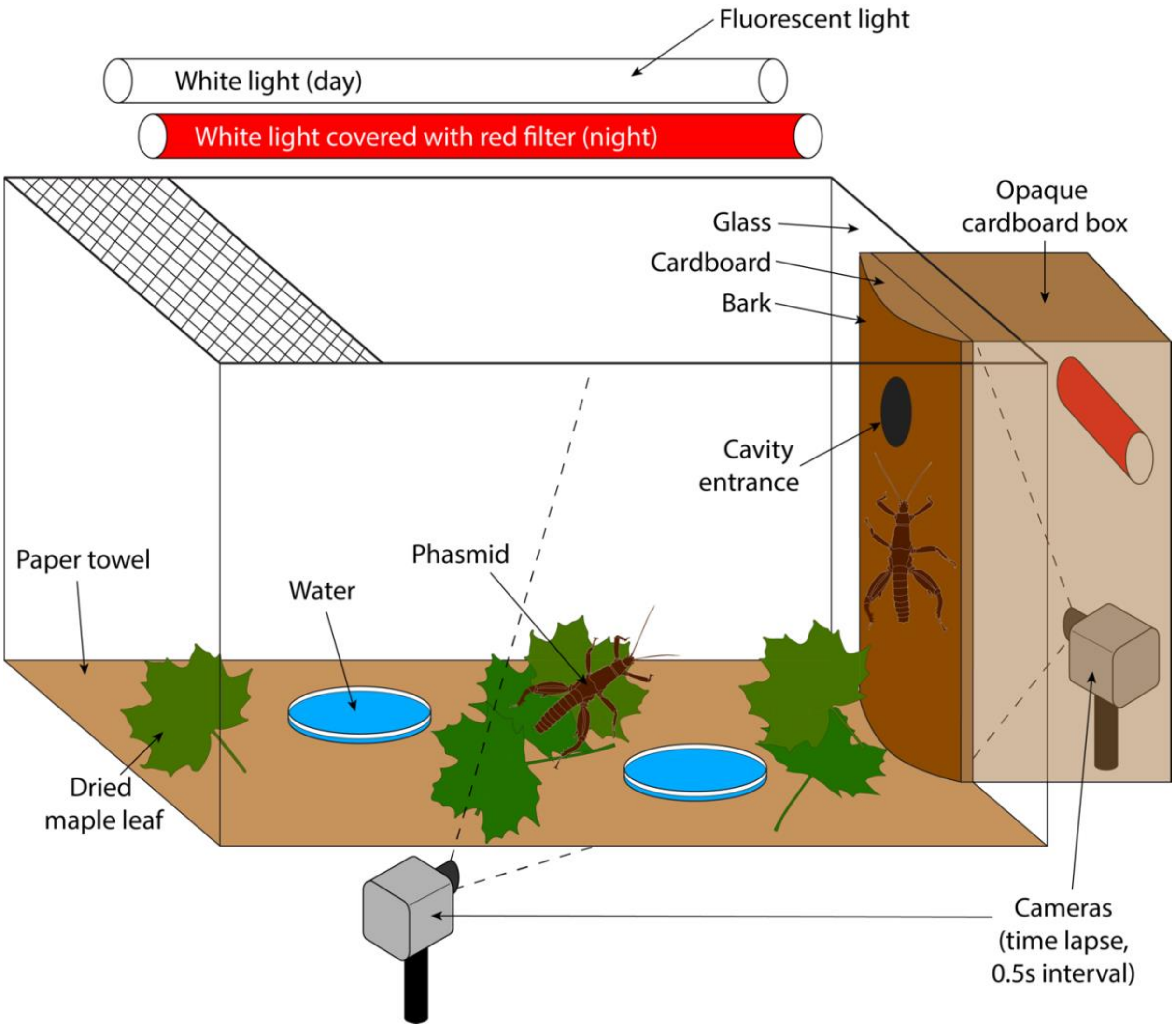


**Figure S2:** Experimental set up for fighting trials.

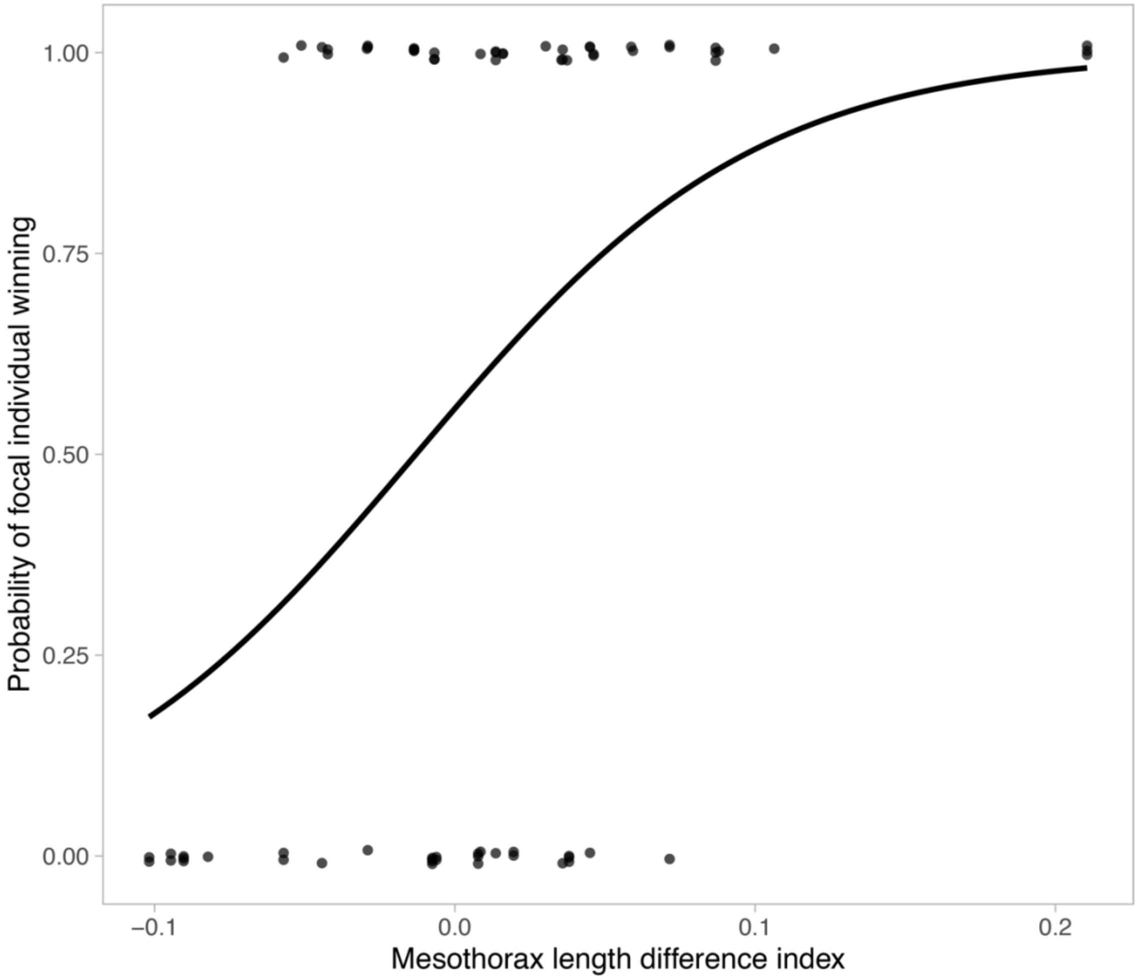


**Figure S3:** Binary GLMM of mesothorax length difference index in randomly matched lab trials against contest outcome inside the daily roosting cavity.

**Datasets S1**: Dataset_Eurycantha_fights_final.xlsx

Measurements_gen1:
Morphological measurements of individuals from the first generation of lab insects (used in randomly matched contests)

- ID: individual ID
- sex: F: female, M: male
- instar: developmental stage. 7: adult.
- date: date of measurement
- body_length: length from head to end of 9th abdominal segment (included) (mm)
- mesothorax_length: length of mesonotum (mm)
- prothorax_width: middle width of pronotum (mm)
- mesothorax_width: middle width of mesonotum (mm)
- head_length: length of head (mm)
- head_width: width of head behind eyes (mm)
- ovipositor_length: length of the female epiproct (mm)
- front_femur_length: length of right front femur (mm)
- front_tibia_length: length of right front femur (mm)
- hind_femur_length: length of right hind femur (mm)
- hind_tibia_length: length of right hind tibia (mm)
- hind_femur_width: width of right hind femur in lateral view at base of largest spine (mm)
- spine_length: length of the largest spine on the right hind femur (mm)
- HL_area: area of the right hind femur measured directly by drawing the outline (mm2)
- spine_area: area of the largest spine on the right hind femur measured directly by drawing the outline (mm2)

Out_fight_outcomes_gen1_wide
Outcomes of randomly matched lab contests (wide format), including only contests outside the cavity

- generation: generation of animals
- fight_ID: Unique ID number of contest
- pair_ID: Unique ID number of male pair
- Trial: Trial ID number
- Container: contest arena ID
- Day: Day when contest happened relative to trial start
- When: when contest happened (day or night)
- Where: when contest happened (inside/outside the cavity)
- Focal: ID of the focal male
- opponent: ID of the opponent male
- first_encounter: is it the first fight between the two males? (1: yes, 0: no)
- ownership: ID of resident male (1: focal, 0: intruder)
- focal_wins: whether focal male wins the contest (1: yes, 0: no)
- mass_focal_before: body mass of focal male before trial (g)
- mass_opponent_before: body mass of opponent male before trial (g)
- mass_focal_after: body mass of focal male after trial (g)
- mass_opponent_after: body mass of opponent male after trial (g)
- body_mass_focal: average body mass of focal (g)
- body_mass_opponent: average body mass of opponent (g)
- mesothorax_length_focal: length of mesonotum (mm) of focal male
- mesothorax_length_opponent: length of mesonotum (mm) of opponent male
- HL_area_focal: hind right femur area ($mm^2$) of focal male
- HL_area_opponent: hind right femur area ($mm^2$) of opponent male
- winner: ID of winner
- loser: ID of loser
- mesothorax_length_winner: length of mesonotum (mm) of winner
- mesothorax_length_loser: length of mesonotum (mm) of loser

- HL_area_winner: hind right femur area ($mm^2$) of winner
- HL_area_loser: hind right femur area ($mm^2$) of loser
- body_mass_winner: average body mass of winner (g)
- body_mass_loser: average body mass of loser (g)
- max_escalation: maximum escalation level/stage reached
- escalation_score: contest cost metric

<u>Out_fight_outcomes_gen1_long</u>
Outcomes of randomly matched lab contests (long format), including only contests outside the cavity
- fight_ID: Unique ID number of contest
- pair_ID: Unique ID number of male pair
- Trial: Trial ID number
- Contestant: ID of contestant
- Outcome: whether contestant won the contest (1: yes, 0: no)
- mesothorax_length: length of mesonotum (mm) of contestant
- HL_area: hind right femur area ($mm^2$) of contestant
- body_mass: average body mass of contestant (g)
- day: Day when contest happened relative to trial start
- first_encounter: is it the first fight between the two males? (1: yes, 0: no)
- ownership: is the contestant the resident male? (1: yes, 0: no)

<u>In_dominance_outcomes_gen1_wide</u>
Outcomes of randomly matched lab contests (wide format), including only contests inside the cavity
- fight_ID: Unique ID number of contest
- pair_ID: Unique ID number of male pair
- Trial: Trial ID number
- Day: Day when contest happened relative to trial start
- Focal: ID of the focal male
- opponent: ID of the opponent male
- focal_wins: whether focal male wins the contest (1: yes, 0: no)
- body_mass_focal: average body mass of focal (g)
- body_mass_opponent: average body mass of opponent (g)
- mesothorax_length_focal: length of mesonotum (mm) of focal male
- mesothorax_length_opponent: length of mesonotum (mm) of opponent male
- HL_area_focal: hind right femur area ($mm^2$) of focal male
- HL_area_opponent: hind right femur area ($mm^2$) of opponent male

<u>In_dominance_outcomes_gen1_long</u>
Outcomes of randomly matched lab contests (long format), including only contests inside the cavity
- fight_ID: Unique ID number of contest
- pair_ID: Unique ID number of male pair
- Trial: Trial ID number
- Contestant: ID of contestant
- Outcome: whether contestant won the contest (1: yes, 0: no)
- mesothorax_length: length of mesonotum (mm) of contestant
- HL_area: hind right femur area ($mm^2$) of contestant
- body_mass: average body mass of contestant (g)

<u>Repeated_trials_gen1</u>
Outcomes of contests involving the smallest male (loser) in the randomly matched lab trials (outside the cavity)
- Trial: Trial ID number
- fight_ID: Unique ID number of contest
- focal_smallest: ID of focal loser
- opponent_larger: ID of winner

- focal_mesothorax_length: length of mesonotum (mm) of focal male
- opponent_mesothorax_length: length of mesonotum (mm) of opponent male
- escalation_score: contest cost metric

Fighting_mating_success_gen1
Mating success data of males involved in the randomly matched lab trials
- Trial: Trial ID number
- male_ID: ID of contestant
- fighting_success_inside: fighting rank inside cavity (1: most dominant male, 3: least dominant male)
- fighting_success_outside: fighting rank outside cavity (1: most dominant male, 3: least dominant male)
- fighting_success: average fighting rank
- absolute_copulation_success: number copulation during entire trial
- relative_copulation_success: number copulation during entire trial relative to average.

Mating latency_gen1
Mating latency and copulation duration data of males and females involved in the randomly matched lab trials
- trial: Trial ID number
- day: Day when contest happened relative to trial start
- copulation_ID: unique ID of copulation event
- male_ID: ID of male
- female_ID: ID of female
- female_mating_status: Female mating status (virgin or mated)
- where: where copulation happened in the arena
- when: when copulation happened (day or night)
- copulation_duration: duration of copulation (s)
- latency_to_mate: time between first contact between male and female and onset of copulation
- mesothorax_length_male: length of mesonotum (mm) of male
- mesothorax_length_female: length of mesonotum (mm) of female
- HL_area_male: hind right femur area ($mm^2$) of male
- HL_area_female: hind right femur area ($mm^2$) of female

Female_fecundity_mating_gen1
Fitness data of mated and parthenogenetic females
- female_ID: ID of female
- mating_status: female mating treatment (P: parthenogenesis, M: mated)
- lifetime: adult longevity from final molt to death (days)
- delay_first_egg: Time between female final molt and laying of the first egg (days)
- total_eggs_laid: Total lifetime number of eggs laid by the female
- total_egg_mass: Total cumulative mass of eggs laid by the female during its lifetime (g)
- single_egg_mass: average mass of single egg (g)
- hatchling_number: number of hatchlings
- hatching_rate: egg hatching rate
- mesothorax_length_female: length of mesonotum (mm) of female
- incubation_duration: egg development duration, from laying of first egg to first hatching.

Offspring_survival_gen1
Offspring survival data of mated and parthenogenetic females
- box: ID of offspring cage
- mating_status: mating treatment of female (M: mated, P: parthenogenetic)
- total_number: total number of hatchlings in the cage
- L1_deaths: number of hatchlings found dead in the cage before reaching the second instar
- death_rate: number of dead hatchlings/total number of hatchlings in the cage

Measurements_gen2

Morphological measurements of individuals from the second generation of lab insects (used in size-matched contests)

- ID: individual ID
- sex: F: female, M: male
- date: date of measurement
- body_length: length from head to end of 9th abdominal segment (included) (mm)
- mesothorax_length: length of mesonotum (mm)
- mesothorax_width: middle width of mesonotum (mm)
- HL_area: area of the right hind femur measured directly by drawing the outline (mm2)
- spine_area: area of the largest spine on the right hind femur measured directly by drawing the outline ($mm^2$)

Out_fight_outcomes_gen2_wide

Outcomes of size-matched lab contests (wide format), including only contests outside the cavity

- generation: generation of animals
- fight_ID: Unique ID number of contest
- pair_ID: Unique ID number of male pair
- Trial: Trial ID number
- Container: contest arena ID
- Day: Day when contest happened relative to trial start
- When: when contest happened (day or night)
- Where: when contest happened (inside/outside the cavity)
- Focal: ID of the focal male
- opponent: ID of the opponent male
- first_encounter: is it the first fight between the two males? (1: yes, 0: no)
- ownership: ID of resident male (1: focal, 0: intruder)
- focal_wins: whether focal male wins the contest (1: yes, 0: no)
- body_mass_focal: average body mass of focal (g)
- body_mass_opponent: average body mass of opponent (g)
- body_mass_female: average body mass of female (g)
- mesothorax_length_focal: length of mesonotum (mm) of focal male
- mesothorax_length_opponent: length of mesonotum (mm) of opponent male
- HL_area_focal: hind right femur area ($mm^2$) of focal male
- HL_area_opponent: hind right femur area ($mm^2$) of opponent male
- size_matched: whether the contest is size matched (mesothorax length of contestants do not differ by more than 5%).
- winner: ID of winner
- loser: ID of loser
- mesothorax_length_winner: length of mesonotum (mm) of winner
- mesothorax_length_loser: length of mesonotum (mm) of loser
- HL_area_winner: hind right femur area ($mm^2$) of winner
- HL_area_loser: hind right femur area ($mm^2$) of loser
- body_mass_winner: average body mass of winner (g)
- body_mass_loser: average body mass of loser (g)
- max_escalation: maximum escalation level/stage reached
- escalation_score: contest cost metric

Out_fight_outcomes_gen2_long

Outcomes of size-matched lab contests (long format), including only contests outside the cavity

- fight_ID: Unique ID number of contest
- pair_ID: Unique ID number of male pair
- Trial: Trial ID number
- Contestant: ID of contestant

- Outcome: whether contestant won the contest (1: yes, 0: no)
- mesothorax_length: length of mesonotum (mm) of contestant
- HL_area: hind right femur area ($mm^2$) of contestant
- body_mass: average body mass of contestant (g)
- day: Day when contest happened relative to trial start
- first_encounter: is it the first fight between the two males? (1: yes, 0: no)
- ownership: is the contestant the resident male? (1: yes, 0: no)

Field_fight_outcomes_PNG_wide
Outcomes of contests observed in the field (wide format)
- fight_ID: Unique ID number of contest
- night: date of observation
- pair_ID: Unique ID number of male pair
- Focal: ID of the focal male
- opponent: ID of the opponent male
- focal_wins: whether focal male wins the contest (1: yes, 0: no)
- ownership: ID of resident male (1: focal, 0: intruder)
- female_around: is a female close to the fighting males?
- description: qualitative description of the contest
- pixel_length_focal: body length of focal male in pixels
- pixel_length_opponent: body length of opponent male in pixels

Fights_behavioral_sequence_gen1 (randomly matched lab contests)
Fights_behavioral_sequence_gen2 (size matched lab contests)
Fights_behavioral_sequence_PNG (field contests)
Behavioral transitions that occurred during randomly matched lab contests, size matched lab contests and field contests respectively
- trial: Unique ID number of the trial
- fight_ID: Unique ID number of contest
- behavior1: First behavior of a transition
- behavior2: Second behavior of a transition
